# Distinct modes of N-terminal remodeling through a conserved inhibitor pocket in connexin-46/50

**DOI:** 10.64898/2026.10.26.762086

**Authors:** Joshua M. Jarodsky, Janette B. Myers, Steve L. Reichow

## Abstract

Selective pharmacological control of gap junction communication remains challenging because existing small-molecule connexin inhibitors act across multiple isoforms. The structural basis of this broad activity remains incompletely understood. Here, we used single-particle cryo-electron microscopy to examine native connexin-46/50 gap junction channels treated with two commonly used and chemically distinct inhibitors, 2-aminoethoxydiphenyl borate (2-APB) and carbenoxolone (CBX). Both treatments perturb the hydrophobic N-terminal (NT)-docking groove, but produce distinct conformational ensembles. 2-APB yields ordered destabilized-open and gated NT states resembling conformations observed during physiological pH- and Ca^2+^-dependent gating, with ligand- and lipid-associated densities accompanying the gated state. In contrast, CBX-compatible density occupies the same groove and is associated with loss of an ordered NT conformation, consistent with steric competition and altered physicochemical complementarity at the NT-docking site. Comparison with inhibitor-treated connexin-32 and connexin-36 structures identifies this region as a structurally conserved, pharmacologically sensitive pocket. These findings reveal how chemically diverse inhibitors perturb a conserved element of the native gating machinery while producing divergent NT-gating ensembles, providing a structural rationale for broad cross-isoform activity and identifying sequence-variable pocket margins that may support more selective modulation.

## Introduction

Gap junction channels, formed by connexin proteins, establish direct pathways for electrical and metabolic communication between neighboring cells, coordinating long-range signaling and tissue homeostasis throughout the body^1,2^. Given their broad physiological roles, connexin dysfunction is associated with diverse diseases, including blindness, deafness, arrhythmia, neuropathy, skin disorders, and cancer^3–5^. Although intercellular coupling is essential for normal tissue function, it can also propagate cytotoxic signals through the bystander effect during pathological conditions such as ischemia, arrhythmia, and seizure^6–13^. Pharmacological control of gap junction communication is therefore valuable both for investigating connexin function and as a potential therapeutic strategy^14^. However, available connexin inhibitors generally lack potency and isoform selectivity, limiting their experimental and therapeutic utility.

Gap junction channels assemble through docking of two hexameric connexin hemichannels from adjacent cells, forming an aqueous pore that facilitates the direct exchange of ions and metabolites^15,16^. Each connexin subunit contains four transmembrane helices (TM1–4), two extracellular loops that mediate hemichannel docking, and an N-terminal (NT) domain that lines the pore and participates in channel gating^17–19^ (**Fig. 1a**). In the conductive open-state of Cx46/50 channels, the hydrophobic face of the NT is docked within a luminal groove formed by TM1/TM2, and TM1 of the neighboring subunit, stabilizing the NT against the channel wall and leaving the pore unobstructed^20–25^. Structural studies of Cx46/50 and other connexin isoforms have shown that physiological gating signals, including pH, Ca^2+^, and CO_2_, remodel the NT conformational ensemble, consistent with playing a central role in the channel gating mechanism^22,26–30^. The surrounding lipid environment can also facilitate or stabilize these rearrangements^22,26,30,31^, highlighting a role for lipids in shaping the conformational landscape of the NT domain.

**Figure 1:**
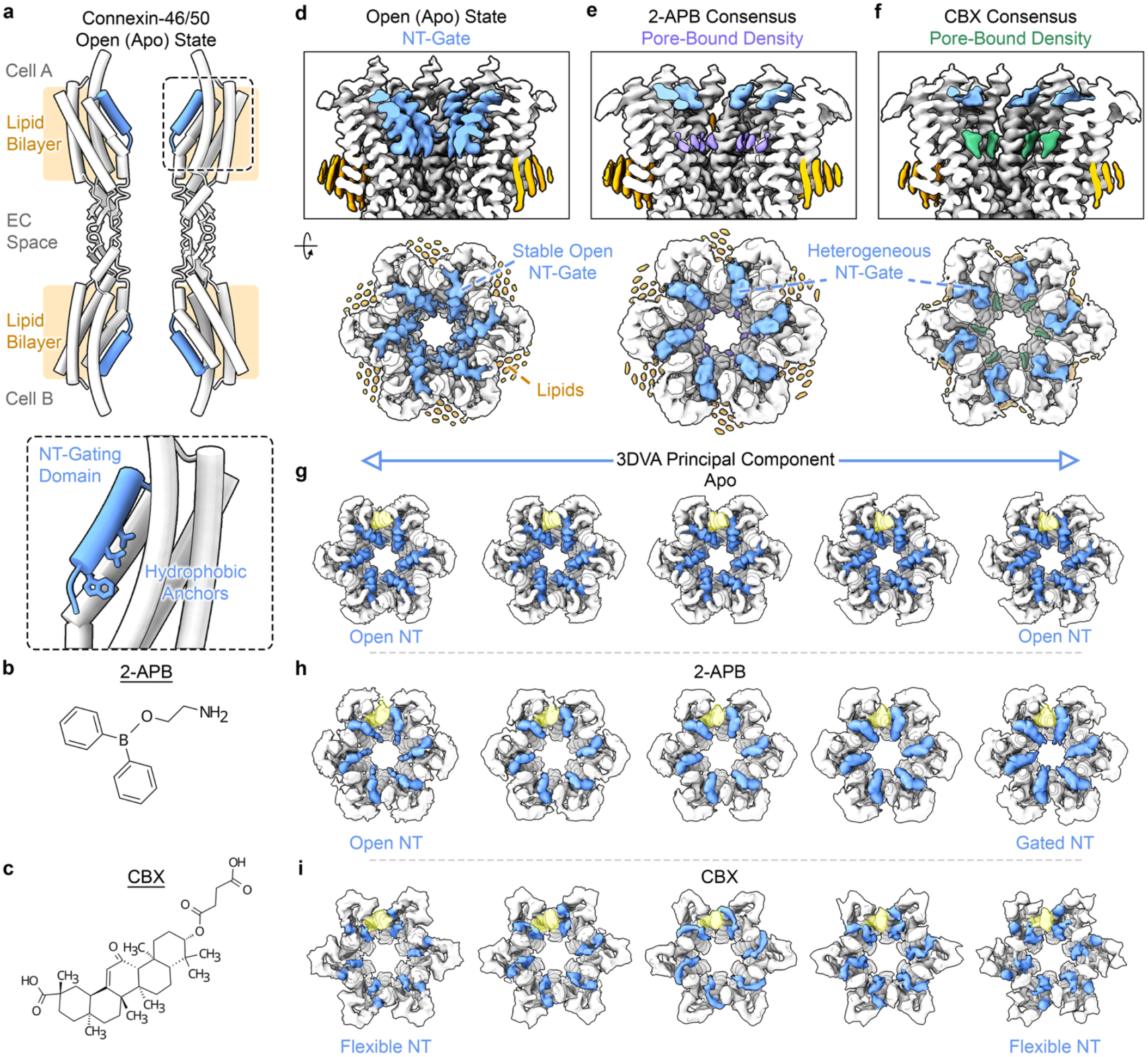
2-APB and CBX induce distinct NT/TM2 conformational ensembles in Cx46/50. **a,** Schematic of a connexin subunit with the N-terminal domain (NT) in its open-state position (PDB 7JKC). *Inset,* close up of the NT-gating domain showing the hydrophobic anchor residues W4, L7 and L11. **b,c,** Chemical structures of (*b*) 2-aminoethoxydiphenyl borate (2-APB) and (*c*) carbenoxolone (CBX). **d–f,** Consensus cryo-EM maps of Cx46/50 in the (*d*) previously determined apo open-state (EMD-73885) and following treatment with (*e*) 2-APB or (*f*) CBX. Maps are shown in cross-sectional (left) and cytoplasmic (right) views. NT density is highlighted in light blue, annular lipid density in orange, 2-APB-associated pore density in purple, and CBX-associated density in green. **g–i,** Representative reconstructions along the principal mode of variability identified by 3D variability analysis (3DVA) for (*g*) apo Cx46/50 (EMPIAR-13131), (*h*) 2-APB-treated Cx46/50, and (*i*) CBX-treated Cx46/50. NT density is highlighted in light blue and the cytoplasmic portion of TM2 in yellow. **g** Under apo conditions, the NT remains near its open-state position with only minor variability in the proximal NT and TM2. **h** Following 2-APB treatment, ordered NT density spans open-like and gated-like conformations accompanied by coordinated inward displacement of TM2. **i** Following CBX treatment, the NT remains largely unresolved throughout the component, while TM2 exhibits limited variability without the coordinated displacement observed with 2-APB, consistent with delocalized conformational heterogeneity of the NT.

Connexins are inhibited by chemically diverse small molecules, including long-chain alcohols, fatty acids, quinoline derivatives, and glycyrrhetinic acid derivatives^32–34^. Among the most widely used are 2-aminoethoxydiphenyl borate (2-APB; **Fig. 1b**) and carbenoxolone (CBX; **Fig. 1c**), both of which inhibit multiple connexin isoforms^12,33,35–39^ and act on other membrane proteins, including the related pannexins^40,41^, and unrelated targets, such as 11β-hydroxysteroid dehydrogenase, store-operated calcium-entry channels, and transient receptor potential channels^42–44^. Recent structures of Cx32 and Cx36 treated with mefloquine, 2-APB, hexanol, or arachidonic acid revealed assigned ligand densities near TM1 and TM2 associated with a range of ordered and unresolved NT conformations^45–47^. Several structures also contained additional pore-associated densities assigned as lipids, including discrete lipid-like features and broad bilayer-like density proposed to contribute to pore occlusion. However, differences in connexin isoforms, biochemical environment, and density assignment complicate direct comparison among these structures. Moreover, these systems lack the matched series of physiologically induced gating states available for Cx46/50^20–22,30^, limiting interpretation of how inhibitor-associated conformations relate to native gating mechanisms. Consequently, how chemically diverse inhibitors perturb the NT-gating machinery, and which features of their action are shared across connexin isoforms, remain incompletely understood.

Here, we used single-particle cryo-electron microscopy (cryo-EM) to compare the structural effects of the commonly used and chemically distinct inhibitors 2-APB and CBX on native Cx46/50 gap junction channels. Under apo cryo-EM conditions, these channels predominantly adopt a stable open-state conformation in which the NT-gating domain is docked within a hydrophobic groove lining the channel lumen^20–24^, and previously determined low-pH and Ca^2+^-gated structures provide reference states for interpreting inhibitor-associated changes^22,30^. Treatment with 2-APB produces ordered NT conformations that overlap with physiological gating, with gated and open-like states coexisting asymmetrically within individual channels consistent with noncooperative transitions among subunits. CBX produces a distinct response in which the NT-gating domain becomes delocalized. Both responses are associated with ligand-associated densities within the same NT-docking groove. Comparisons with inhibitor-treated Cx32 and Cx36 identify this region as a conserved, pharmacologically sensitive pocket. Together, these findings show how chemically diverse inhibitors can act through a shared element of the native gating machinery while producing distinct conformational responses, clarifying both the basis of broad inhibitor activity across connexin isoforms and identifying potential routes toward improved isoform selectivity.

## Results

### 2-APB and CBX induce distinct changes to the Cx46/50 NT-gating domain

Native Cx46/50 channels were purified from aged ovine lens tissue and reconstituted into MSP1E1 nanodiscs containing dimyristoylphosphatidylcholine (DMPC) under conditions previously shown to stabilize the open-state (EMD-73885; **Fig. 1d; Supplementary Fig. 1**)^21,22,48–50^. Samples were treated with 300 µM 2-APB in 0.5% DMSO (vol vol^−1^), corresponding to ∼10- and 100-fold the reported IC_50_ values for Cx46 and Cx50, respectively^35^, or with 5 mM CBX. Because CBX inhibition has not been quantitatively characterized for Cx46 or Cx50, this concentration was selected based on published IC_50_ values for other connexin isoforms^36,37,51^. Negative-stain EM confirmed that the reconstituted nanodisc complexes remained intact following treatment with either inhibitor (**Supplementary Fig. 1**). Moreover, our previous structural analysis of bovine Cx46/50, which shares >95% sequence identity with ovine Cx46/50 across the structured regions, showed that treatment with 1% DMSO preserved the stabilized open-state conformation under otherwise similar conditions^24^.

Preliminary single-particle cryo-EM analysis produced consensus reconstructions of 2-APB-treated Cx46/50 (Cx46/50_2-APB_) and CBX-treated Cx46/50 (Cx46/50_CBX_) at global resolutions of 2.1 Å and 2.4 Å, respectively (**Fig. 1e,f**). Although the transmembrane and extracellular regions were well resolved in both maps, density corresponding to the NT domain was weak in Cx46/50_2-APB_ and largely absent in Cx46/50_CBX_. This contrasts with the well-defined open-state NT conformation observed in the previously determined apo-state Cx46/50 structure (**Fig. 1d**)^22^. Thus, treatment with either inhibitor was associated with loss of the uniformly ordered open-state NT conformation.

To distinguish the underlying conformational behavior of the NT-gating domain, the Cx46/50_2-APB_ and Cx46/50_CBX_ datasets were independently analyzed by 3D variability analysis (3DVA)^52^ at the gap junction level using symmetry expanded particles and compared with equivalent analysis of our previously published apo-state Cx46/50 dataset (EMPIAR-13131; **Fig. 1g–i** and **Supplementary Movie 1**)^22^. Under apo conditions, the NT remained near its open-state position across the primary mode of variability, accompanied by only minor perturbations of the proximal NT and cytoplasmic portion of TM2 (**Fig. 1g**). In contrast, Cx46/50_2-APB_ sampled a continuum of ordered NT conformations spanning open-like and gated-like states, with NT displacement coupled to movement of the cytoplasmic portion of TM2 (**Fig. 1h**). This coordinated behavior resembles that observed under physiological Cx46/50 gating conditions^22,30^. In Cx46/50_CBX_, however, the NT remained largely unresolved throughout the variability analysis, and no discrete open- or gated-like NT conformation could be recovered (**Fig. 1i**). The cytoplasmic portion of TM2 exhibited only minor movements without the coordinated NT displacement observed following 2-APB treatment. Thus, although both inhibitors perturb the ordered open-state NT conformation, 2-APB and CBX produce distinct NT/TM2 conformational ensembles that are examined in greater detail below.

### 2-APB induces an ensemble of NT-gated states with ligand- and lipid-associated densities

Deep classification of the 2-APB-treated Cx46/50 dataset resolved two D6-symmetrized reconstructions distinguished by their NT-domain conformations, termed the destabilized-open and gated states, which were refined to 2.4 Å and 2.8 Å resolution, respectively (**Fig. 2; Supplementary Figs. 2,3; Supplementary Table 1**). Separate Cx46 and Cx50 atomic models, corresponding to TM1-4, EC1-2 and regions of the NT, could be fit equally well into each reconstruction; however, as in previous studies^20–22,24,30^, the heteromeric and heterotypic arrangement of Cx46 and Cx50 within the native channels could not be determined because of their high (>80%) sequence identity across the structured regions. Cx46 is therefore presented as the archetype, with differences in Cx50 noted where relevant.

**Figure 2:**
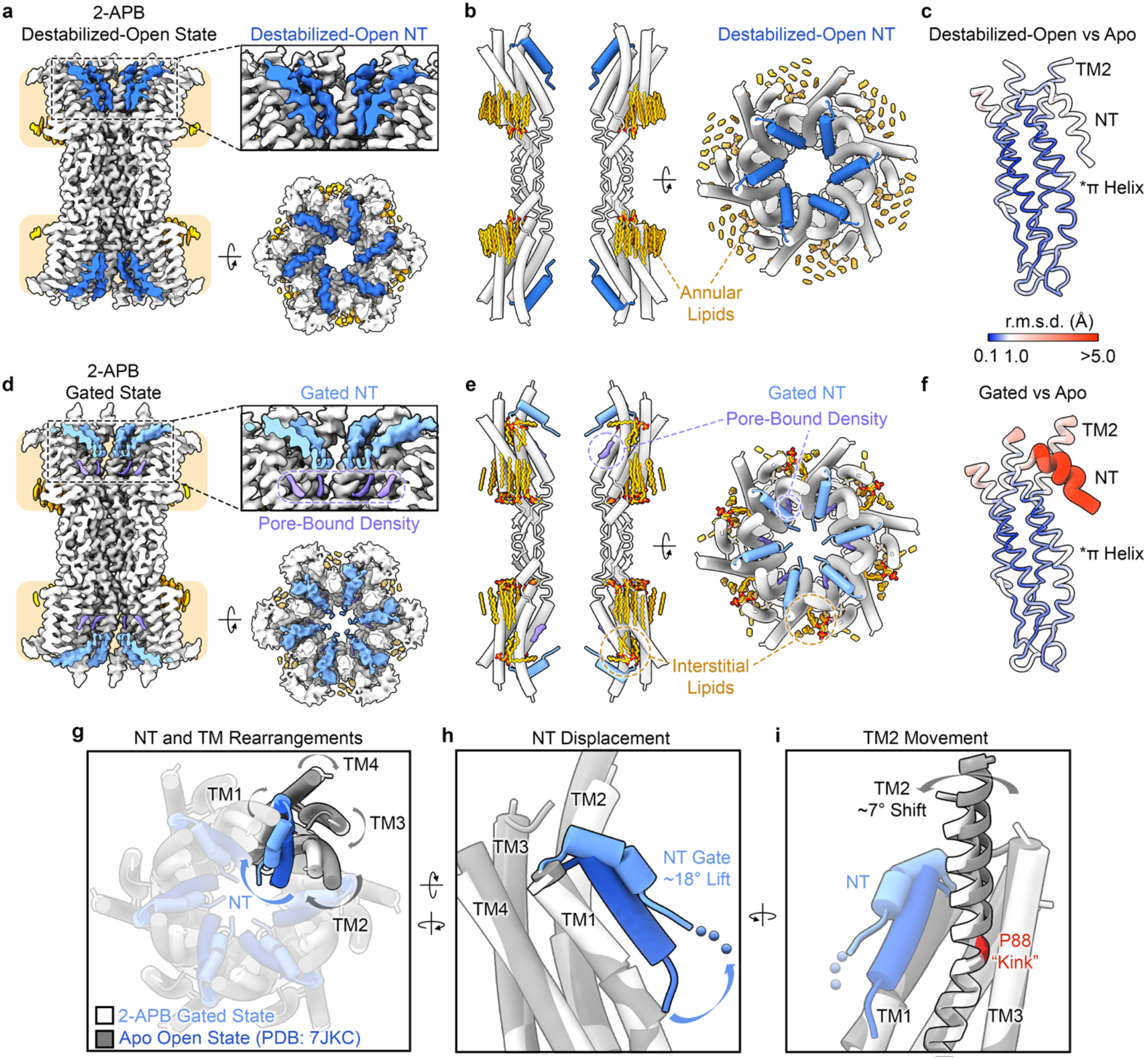
2-APB treatment induces ordered destabilized-open and gated Cx46/50 states. **a**, Cross-sectional (left), close-up of the NT-region (upper right) and cytoplasmic (lower right) views of the cryo-EM map of the 2-APB destabilized-open state of Cx46/50 in DMPC nanodiscs. **b,** Cross-sectional (left) and cytoplasmic (right) views of the corresponding atomic model. The NT domains are shown in dark blue and resolved lipids in orange. **c,** Cα root-mean-square deviation (r.m.s.d.) analysis of the destabilized-open state relative to apo open-state Cx46 (PDB 7JKC). **d,e,** Corresponding views of the (*d*) cryo-EM map and (*e*) atomic model of the 2-APB-gated state, with the NT domains shown in light blue, pore-bound densities in purple, and resolved lipids in orange. **f,** Cα r.m.s.d. analysis of the 2-APB-gated state relative to apo open-state Cx46. In *(c)* and *(f)*, tube thickness and color indicate the magnitude of structural deviation according to the scale shown. **g,** Overlay of Cx46 atomic models from the apo open-state (PDB 7JKC; NT, dark blue; transmembrane helices, dark gray) and 2-APB-gated state (NT, light blue; transmembrane helices, light gray). When viewed from the cytoplasmic side, displacement of the NT toward the pore axis is accompanied by a clockwise rearrangement of the TM1–TM4 bundle. **h,** Close-up of the NT displacement. Relative to the apo open-state, the resolved NT is displaced by ∼18° and 5.3 Å toward the pore axis. The light-blue dashed segment indicates unmodeled residues G2–S5; side chains of residues 6–17 were truncated to Cβ because of limited local resolution. **i,** Close-up of the coupled TM2 movement. The cytoplasmic portion of TM2 tilts by ∼7° around the conserved P88 kink (red), corresponding to a displacement of ∼3.1 Å.

The 2-APB destabilized-open state closely resembles destabilized-open conformations previously observed under low-pH and Ca^2+^-gating conditions^22,30^ (**Fig. 2a,b**). In this state, the NT remains docked within the hydrophobic groove formed by TM1/TM2, but is slightly displaced relative to the stabilized open-state (local Cα r.m.s.d. ≈ 1.2 Å for residues 3–17). This rearrangement is accompanied by a minor displacement of the cytoplasmic portion of TM2 (local Cα r.m.s.d. ≈ 0.8 Å for residues 88–101), consistent with coupling between the NT and TM2 (**Fig. 2c**).

The 2-APB-gated state contains a more pronounced displacement of the NT, and disengagement from the channel lumen toward the center of the pore (**Fig. 2d–i**). This movement is accompanied by a coordinated clockwise rearrangement of the cytoplasmic portions of the transmembrane helices, whereas EC1 and EC2 remain comparatively unchanged (**Fig. 2f,g**). Relative to the apo open-state, the 2-APB-gated state has an overall Cα r.m.s.d. of ∼0.8 Å, with the largest differences localized to the NT and cytoplasmic portion of TM2. The resolved portion of the NT is displaced by ∼18° toward the pore (local Cα r.m.s.d. ≈ 5.2 Å for residues 6–17), while TM2 tilts by ∼7° around the conserved proline kink (P88 in Cx46/50), producing a displacement of ∼3.1 Å at its cytoplasmic end (local Cα r.m.s.d. ≈ 2.0 Å for residues 88–104) (**Fig. 2h,i**). Through these rearrangements, the NT remains coupled to TM2 via contacts between A14 and L93 in Cx46 (V14 and V93 in Cx50). The gated NT is less well resolved than the remainder of the channel: residues G2–S5 could not be modeled, and the side chains of residues 6–17 were truncated to Cβ atoms. Consequently, the structure establishes displacement of the NT toward the pore but does not define the position of the proximal NT residues or the resulting pore constriction. Nonetheless, the overall NT/TM2 conformation closely resembles previously described gated states of Cx46/50^22,30^.

The 2-APB-gated reconstruction also contains non-protein densities at the lateral subunit interfaces and within the NT docking groove (**Figs. 2d,e and Fig. 3a,b**). At the subunit interface near the cytoplasmic leaflet, a splayed density was modeled as an interstitial DMPC lipid (IL; **Fig. 3a–d**). One acyl chain contacts hydrophobic residues along TM2 and TM3, including F84, T87, P88, I/V91, H95, F148/C160, F152/164, and F156/168 (Cx46/50 numbering), whereas the second extends toward the channel lumen and contacts TM2 and TM1 of the neighboring subunit. Additional pore-bound densities occupy the hydrophobic groove formed by TM1/TM2, and TM1 of the neighboring subunit (**Fig. 3e,f**). Under apo conditions, this groove accommodates the conserved NT anchor residues W4, L7, and L11 that stabilize the open-state (**Fig. 3g**). In the 2-APB-gated state, the NT is displaced from this site, and the exposed groove is instead occupied by densities contacting hydrophobic residues along TM1 and TM2. The repositioned NT lies above these pore-bound densities, with the anchoring residues L7 and L11 directed toward the occupied groove.

**Figure 3:**
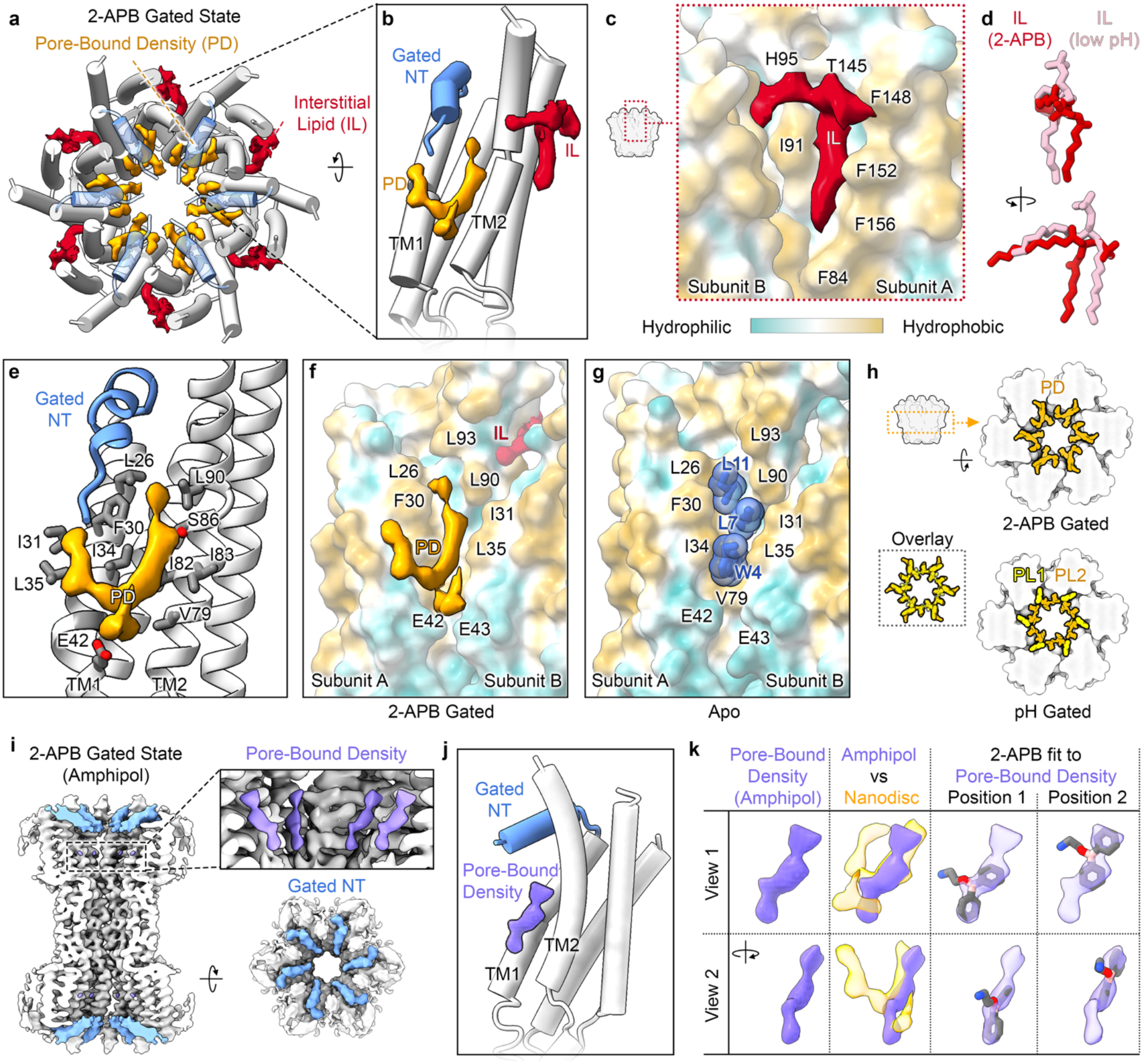
Ligand- and lipid-associated densities occupy the NT-docking pocket in the 2-APB-gated state. **a,b**, Atomic model of the 2-APB-gated Cx46/50 channel shown in (*a*) cytoplasmic view and (*b*) close-up view of a Cx46 subunit. NT domains are light blue, transmembrane helices gray, the segmented interstitial lipid (IL) red and pore-bound density orange. **c,** IL shown at the hydrophobicity-colored subunit interface viewed from the cytoplasmic leaflet. One acyl chain contacts TM2 and TM3, whereas the other extends into the pore and contacts TM1 of the neighboring subunit. **d,** Comparison of the ILs modeled in the 2-APB- and low-pH-gated states (PDB 9Z9X), showing similar positions but subtle differences in one acyl chain and the headgroup. The 2-APB-associated IL headgroup was modeled only through the phosphate. **e,** Pore-bound density occupying the hydrophobic TM1/TM2 groove beneath the displaced NT. Contacting residues are shown as dark-gray sticks. **f,g,** Comparison of the (*f*) 2-APB-gated state, in which pore-bound density occupies the NT-docking site and (*g*) apo open-state (PDB 7JKC), in which conserved hydrophobic residues W4, L7 and L11 anchor the NT within the groove. **h,** Cross-sectional views comparing the pore-bound densities in the 2-APB-gated state with pore-bound lipids PL1 and PL2 in the low-pH gated state (EMD-73962). The densities occupy overlapping regions but exhibit distinct packing arrangements. **i,** Cross-sectional (left) and cytoplasmic (right) views of 2-APB-treated Cx46/50 in amphipol, with NT and pore-bound densities highlighted in light blue and purple, respectively. Inset, shows the locally filtered pore-bound densities. **j,** Cx46 subunit model with the segmented pore-bound density observed in amphipol. **k,** Two orthogonal views comparing the pore-bound densities resolved in amphipol (purple) and nanodiscs (orange). Two candidate 2-APB poses fit within the pore-bound density observed in amphipol, which partially overlaps the density observed in nanodiscs.

Similar interstitial and pore-bound densities were previously resolved in the low-pH-gated state of Cx46/50 and shown to be required for the gating mechanism^22^. The interstitial lipids occupy comparable sites at the lateral subunit interface, although their acyl-chain and headgroup positions differ (**Fig. 3d**). Likewise, although not identical, the 2-APB-associated pore densities overlap with the PL1 and PL2 densities modeled in the low-pH gated structure, which were proposed to form a hydrophobic gasket beneath the displaced NT (**Fig. 3h**). Thus, 2-APB and low pH are associated with similar NT/TM2 rearrangements and occupation of the exposed NT docking region by lipid-like densities. In the 2-APB-treated nanodisc structure, however, the limited resolvability of these densities and the possibility that 2-APB occupies the same region precluded their unambiguous assignment.

To help distinguish inhibitor-from lipid-associated density and potential lipid-dependency, Cx46/50 was reconstituted into amphipol A8-35 without added phospholipid, under conditions previously shown to stabilize the open state^20^, and then treated with 300 µM 2-APB in 0.5% DMSO (vol vol^-1^). Single-particle cryo-EM analysis resolved a gated conformation at 3.2 Å resolution (**Fig. 3i; Supplementary Figs. 1, 4, 5; Supplementary Table 1**). No discrete open-state class was recovered from this dataset. The amphipol structure closely resembles the 2-APB-gated state in nanodiscs (global Cα r.m.s.d. ≈ 0.5 Å) and retains density within the TM1/TM2 hydrophobic groove (**Fig. 3i,j**). These findings demonstrate that a structurally similar gated state and part of the associated pore density can be recovered in the absence of a lipid bilayer environment.

The pore density in the amphipol sample overlaps with a portion of the more extensive pore-bound density observed in nanodiscs, identifying this shared region as the strongest candidate for 2-APB binding (**Fig. 3k**). The diphenyl groups of 2-APB can be accommodated in two positions within the density, but support for the 2-aminoethoxy group is weak, precluding assignment of a unique binding pose. This diffuse density could reflect partial occupancy of multiple poses, binding of the 2-APB hydrolysis product diphenylborinic acid^53^, or averaging of Cx46- and Cx50-associated features. Together, the amphipol and nanodisc structures suggest that the pore-bound density in the 2-APB-treated nanodisc sample contains heterogeneous contributions from 2-APB-related species and phospholipid. Their colocalization beneath the displaced NT is consistent with a model in which inhibitor occupancy competes with open-state NT docking, while lipid occupancy may further stabilize the ordered gated conformation.

To determine whether the destabilized-open and gated NT conformations represented homogeneous channel classes or coexisted within individual channels, we next examined NT heterogeneity at the subunit level. Following D6 symmetry expansion and signal subtraction of the consensus particle stack, individual subunit densities were isolated and subjected to subunit-focused 3DVA (**Supplementary Figs. 2 and 6; Methods**). The primary principal component sampled open and gated NT conformations, with ∼60% of subunits assigned to the open state and 40% to the gated state (**Supplementary Fig. 6**).

Mapping these subunit assignments back to their parent particles showed that most gap junctions contained mixtures of open and gated NT conformations. Fully open and fully gated gap junctions accounted for 0.2% and <0.1% of particles, respectively, whereas particles containing four or five gated subunits were most abundant, each accounting for ∼22% of the population (**Supplementary Fig. 6**). At the hemichannel level, two gated subunits represented the most abundant configuration, and the relative frequencies of spatial arrangements containing two to four gated subunits closely matched those expected from random occupancy (**Supplementary Fig. 6**). Thus, 2-APB treatment produces asymmetric NT ensembles consistent with independent subunit occupancy under the cryo-EM conditions, paralleling the noncooperative gating mechanism proposed for low-pH gating of Cx46/50^22^ and functional evidence that individual subunits can initiate channel gating^54^. Accordingly, the D6-symmetrized maps and models represent idealized, symmetry-averaged views of the destabilized-open and gated subunit conformations and do not imply that all 12 subunits within an individual channel adopt the same state.

### CBX treatment delocalizes the NT-gating domain

Despite extensive heterogeneity analysis, the CBX-treated Cx46/50 dataset yielded one well-resolved structural class, termed the flexible-NT (CBX-flex) state, which was refined to 2.8 Å global resolution (**Fig. 4a–d; Supplementary Figs. 7,8; Supplementary Table 1**). NT residues 2–17 were unresolved and the atomic model was therefore built from S18 onward. Neither 3DVA nor classification using alternative symmetry operators recovered a discrete open- or gated-like NT conformation, consistent with delocalized conformational heterogeneity or flexibility in this region. The remaining transmembrane and extracellular domains closely resemble the apo open-state (global Cα r.m.s.d. ≈ 0.6 Å), with the largest difference localized to the cytoplasmic portion of TM1, which tilts by ∼5.5° away from the lumen and toward the lipid bilayer (**Fig. 4e**). By contrast, the cytoplasmic portion of TM2 undergoes minimal displacement, suggesting an uncoupling of the NT/TM2 rearrangement observed following 2-APB treatment.

**Figure 4:**
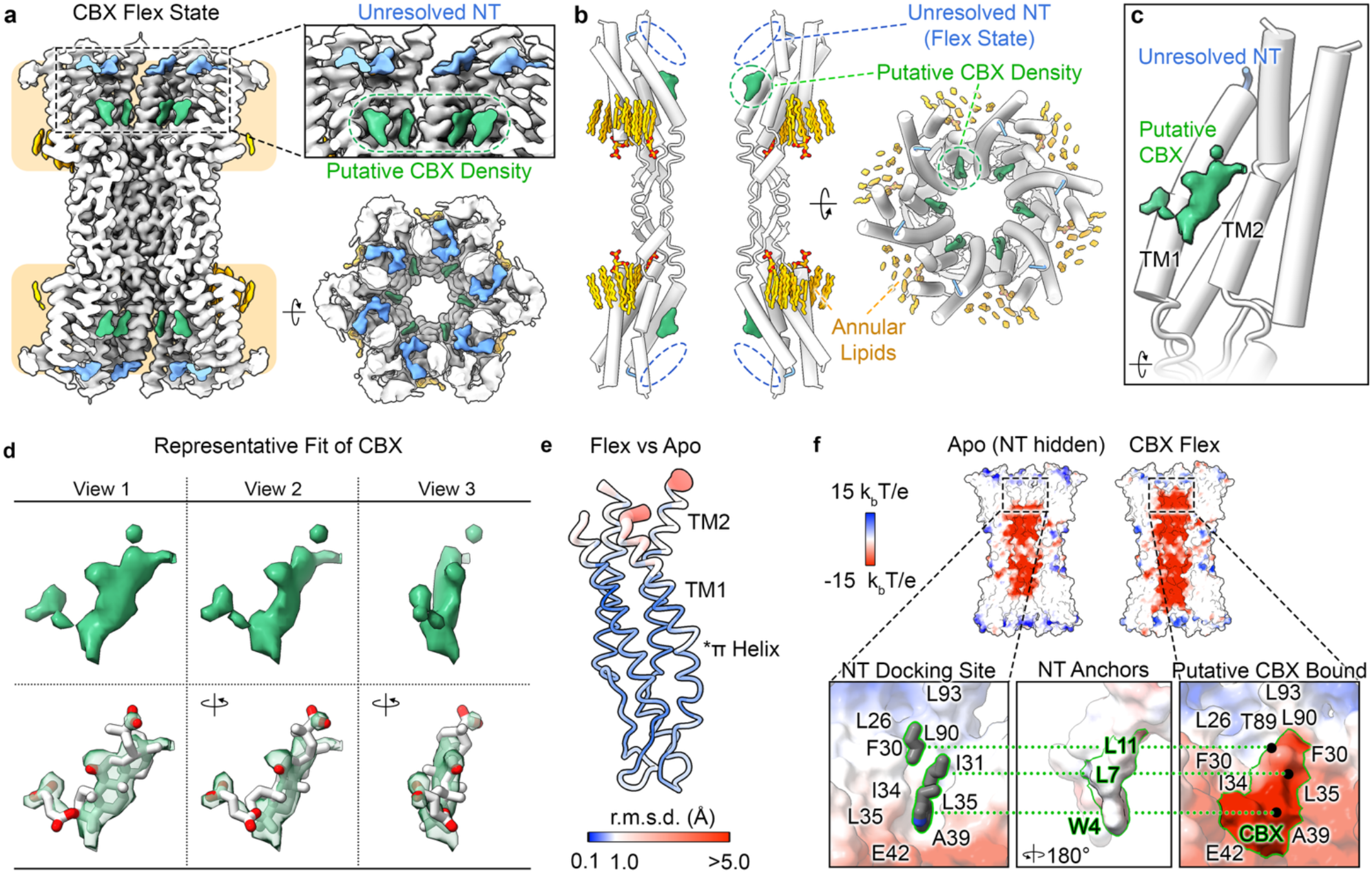
CBX treatment induces delocalized conformational heterogeneity of the NT. **a,b**, Cross-sectional (left), close-up of the NT-region (upper right) and cytoplasmic (lower right) views of the (*a*) cryo-EM map and (*b*) atomic model of the Cx46/50 CBX-flex state. NT density is shown in light blue, annular lipids in orange and CBX-compatible density in green. NT residues 2–17 were not modeled because corresponding density was unresolved, presumably due to conformational flexibility. **c,** Subunit view of the CBX-flex model with segmented density putatively assigned to CBX shown in green. **d,** Three views of the segmented density (top) and a candidate CBX fit (bottom). **e,** Cα root-mean-square deviation (r.m.s.d.) analysis of CBX-flex Cx46 relative to apo open-state Cx46 (PDB 7JKC). Tube thickness and color indicate the magnitude of deviation (color scale shown). **f,** Cross-sectional views of the Coulombic surface potential of Cx46 in the apo open state with the NT omitted (left) and the CBX-flex state (right) (blue, positive; red, negative; scale shown). Boxed regions indicate the NT-docking pocket. Bottom, enlarged views of the apo NT-docking pocket (left), the conserved hydrophobic NT anchor residues W4, L7 and L11 rotated 180° (center), and a putative CBX pose in the CBX-flex state (right). Green outlines and dotted lines illustrate the spatial correspondence between the open-state NT anchors and the modeled CBX pose. The putative CBX pose overlaps the NT-anchor positions and presents a more negative electrostatic surface within the docking pocket.

Loss of the ordered NT conformation is accompanied by the appearance of non-protein density within the hydrophobic groove formed by TM1/TM2, and TM1 of the neighboring subunit (**Fig. 4a–c**). This density differs in shape from the pore-bound densities resolved in the 2-APB-gated state and is compatible with CBX. Its core accommodates the fused-ring structure of CBX, whereas a weak protrusion is consistent with the hemisuccinate group (**Fig. 4d**). However, at lower contour thresholds, weak extensions on either side of the core density could accommodate the hemisuccinate group in multiple orientations, precluding assignment of a unique binding pose (**Supplementary Fig. 9**). Constrained docking calculations^55^ likewise produced similar scores for the candidate orientations and did not distinguish a preferred pose^55^. The absence of nearby cationic residues that could anchor the negatively charged hemisuccinate group may contribute to this positional heterogeneity.

Regardless of its precise orientation, the CBX-associated density overlaps the position occupied by the hydrophobic NT anchor residues W4, L7, and L11 in the apo open-state, indicating that CBX occupancy would sterically compete with open-state NT docking (**Fig. 4f**). Putative modeling of CBX within this site also alters its calculated Coulombic surface potential. The unoccupied TM1/TM2 groove presents a predominantly neutral, hydrophobic surface, whereas the exposed carbonyl and carboxylate groups of modeled CBX produce a more polar and negatively charged environment. CBX occupancy would therefore disrupt both the steric and physicochemical complementarity of the groove with the hydrophobic face of the amphipathic NT domain. These effects provide a structural model in which well-ordered open and gated NT conformations become inaccessible in the CBX-flex state.

Additional unassigned densities were observed between TM1 and TM2 near the NT hinge and at the membrane-facing subunit interfaces (**Supplementary Fig. 9**). The density near the NT hinge is surrounded by hydrophobic residues but is too small to accommodate CBX. Densities near the lateral subunit interfaces occupy regions similar to the interstitial-lipid site in the 2-APB-gated structure but lack the extended rod-like features characteristic of resolved lipid acyl chains and do not extend into the pore. Relative to the apo and 2-APB-gated states, the CBX-associated displacement of TM1 also narrows the fenestration between neighboring subunits (**Supplementary Fig. 10**). Thus, the CBX-flex state differs from the 2-APB-gated state in both its local protein architecture and its associated non-protein densities, although the identities of the additional densities and their relationship to lipid access remain unresolved.

### Chemically diverse inhibitors converge on a conserved NT-docking pocket across connexin isoforms

Although 2-APB and CBX produce distinct effects on Cx46/50, both perturb the NT-docking region (**Fig. 5a**). To determine whether this region represents a broader site of pharmacological perturbation across the connexin family, we examined its sequence conservation and compared the 2-APB- and CBX-associated densities in Cx46/50 with assigned densities resolved in inhibitor-treated Cx32 and Cx36 structures^45–47^. Superposition of densities associated with 2-APB, CBX, mefloquine, hexanol, and arachidonic acid revealed overlapping footprints within the same luminal TM1/2 pocket, which is lined by conserved hydrophobic residues (**Fig. 5b-e**). Although the resolved details and shapes of these densities vary among the reported structures, their spatial overlap identifies the NT-docking groove as a common site of pharmacological perturbation across connexin isoforms.

**Figure 5:**
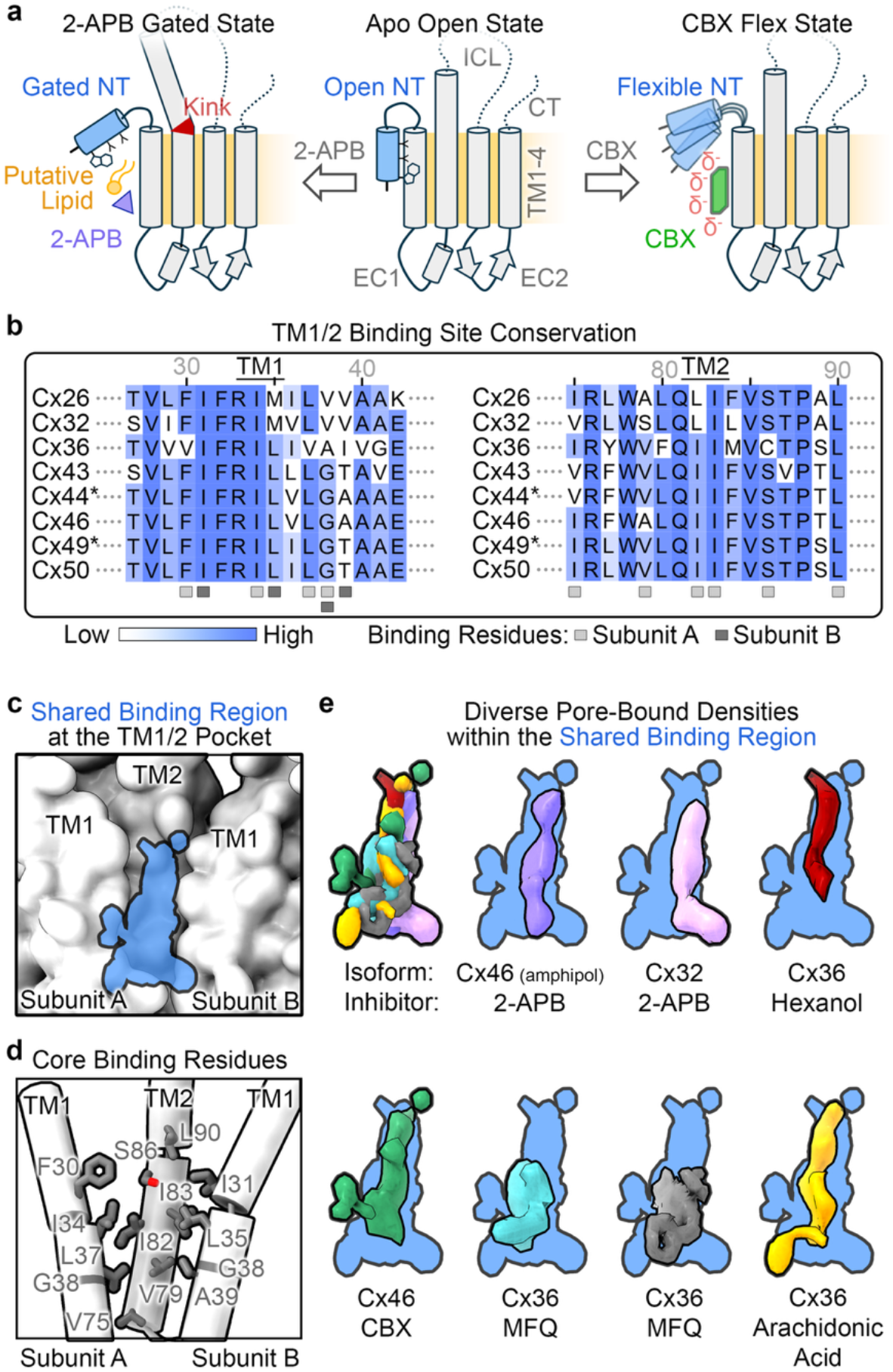
Chemically diverse inhibitors converge on a conserved NT-docking pocket but produce divergent NT states. **a**, Schematic comparing the (center) apo open, (left) 2-APB-gated and (right) CBX-flex states. In the apo state, conserved hydrophobic residues W4, L7 and L11 anchor the NT (blue) within the luminal groove formed by TM1/TM2 and TM1 of the neighboring subunit. In the 2-APB-gated state, putative ligand- and lipid-associated pore densities occupy this groove and accompany inward NT displacement and coupled TM2 movement around the conserved proline kink. A lipid bilayer was not required to recover the gated conformation, although pore-bound lipids may contribute to its stabilization in nanodiscs. The accompanying interstitial lipid observed in nanodiscs is not shown. In the CBX-flex state, CBX-compatible density occupies the same groove and alters the local surface potential from neutral to negative, coinciding with loss of an ordered NT conformation. **b,** Sequence alignment of representative human and ovine connexins (indicated by asterisks) spanning regions of TM1 and TM2. Residues forming the shared pocket are marked below the alignment for subunit A (light gray) and subunit B (dark gray). Residue numbering corresponds to ovine Cx46, and the degree of sequence conservation is indicated by color (scale shown). **c,** Cx46 surface representation showing the shared luminal region occupied by superimposed pore-bound densities (blue) from inhibitor-treated Cx32, Cx36 and Cx46/50 structures. **d,** Cx46 model showing residues forming the shared core of the luminal pocket as labeled gray sticks. **e,** Individual pore-bound densities superimposed on the shared region (blue): 2-APB-treated Cx46/50 in amphipol (purple), CBX-treated Cx46/50 (green), 2-APB-treated Cx32 (pink, site M; EMD-18446), hexanol-treated Cx36 (red; EMD-38322), mefloquine-treated Cx36 (teal; EMD-38327, gray; EMD-18540) and arachidonic-acid-treated Cx36 (yellow; EMD-38320).

The shared core of this pocket is formed predominantly by residues with conserved hydrophobic character contributed by adjacent subunits, including F30, I34, L37, and G38 on TM1; V75, V79, I82, I83, S86, and L90 on TM2; and I31, L35, G38, and A39 on TM1 of the neighboring subunit (Cx46 numbering; **Fig. 5b,d**). Although the overall architecture and hydrophobic character of the pocket are conserved, it also contains sequence-variable positions. A39 in Cx46, for example, corresponds to T39 in Cx50 and may contribute to their differential sensitivity to 2-APB^35^. At the equivalent position in Cx32, substitution of V37 with tryptophan to sterically occlude the pocket produced functional channels with markedly reduced sensitivity to both mefloquine and 2-APB^45^, supporting the pharmacological relevance of this region. The structural conservation and shallow, predominantly hydrophobic character of the pocket therefore provide a rationale for the activity of chemically diverse small-molecule inhibitors across connexin isoforms.

Despite this spatial convergence, inhibitor-treated connexins do not adopt a single conserved NT conformation or mode of pore occlusion. In both Cx46/50 and Cx32, 2-APB treatment produced an ordered displacement of the NT accompanied by pore-bound lipid densities, but the resulting conformations and proposed inhibitor-binding sites differ. In Cx32, the NT adopts a more lifted conformation resembling that of the apo Cx32 hemichannel^56^, and 2-APB was assigned primarily to a distinct site between neighboring NT helices (termed site A)^45^. Weaker density was also observed at a pore-lining position (termed site M), corresponding to the shared TM1/TM2 pocket, together with additional pore-bound densities assigned as lipids. In nanodisc-reconstituted Cx36, mefloquine, arachidonic acid, and hexanol shifted the NT ensemble toward a flexible state accompanied by broad, bilayer-like lipid density proposed to occlude the pore^46^. No comparable bilayer-like occlusion is observed in the 2-APB- or CBX-treated Cx46/50 nanodisc structures. In a separate detergent-based study of Cx36, mefloquine was instead modeled directly within the shared TM1/TM2 pocket^47^. The NT remained unresolved in this structure, but was also unresolved in the corresponding apo state, precluding assignment of an inhibitor-induced NT rearrangement. Together, these comparisons identify the conserved NT-docking groove as a pharmacologically sensitive pocket through which direct inhibitor occupancy, inhibitor-associated lipid remodeling, or both can produce distinct NT conformational ensembles and modes of channel inhibition.

## Discussion

### Distinct structural responses from a shared NT-docking pocket

A particular advantage of the Cx46/50 system is the availability of structurally defined apo-open, low-pH-gated, and Ca²⁺-gated ensembles obtained from the same native channel preparation. This reference landscape shows that the 2-APB-associated states overlap with endogenous gating conformations, whereas CBX produces a distinct loss of NT order. The two inhibitors therefore act on a native gating machinery but redirect its conformational ensemble in different ways (**Fig. 5a**).

Recovery of the 2-APB-gated state in amphipol indicates that a lipid bilayer environment is not strictly required to form this conformation, whereas the more extensive pore-bound densities in nanodiscs suggest that phospholipids may further stabilize or reshape the gated ensemble. By contrast, CBX-compatible density sterically overlaps the NT anchor site and, in the modeled poses, introduces a more polar and negatively charged surface into the otherwise neutral, hydrophobic groove. CBX occupancy could therefore disrupt stable NT docking without favoring a single alternative conformation. The associated narrowing of the intersubunit fenestration could additionally restrict lipid access, although the relationship between fenestration width and lipid occupancy remains speculative. Functional studies support channel closure under saturating CBX conditions, but the unresolved NT density prevents identification of a specific structural constriction. The CBX-flex structure may therefore represent a conformationally heterogeneous nonconducting ensemble rather than a single, well-defined occluded state.

Comparisons with inhibitor-treated Cx32 and Cx36 reinforce that pharmacological perturbation of the NT-docking region is not coupled to a single structural outcome^45–47^. Depending on the connexin, inhibitor, and experimental environment, occupation of this region is associated with ordered or flexible NT conformations, direct inhibitor occupancy, and distinct forms of lipid remodeling. Notably, the broad bilayer-like pore density reported for inhibitor-treated Cx36 in nanodiscs is absent from the Cx46/50 nanodisc structures, indicating that nanodisc reconstitution does not invariably produce this feature. Cross-study comparisons are also complicated by the use of cholesteryl hemisuccinate as an additive, which is shown to influence NT conformation and pore-lipid occupancy in Cx32 and Cx43^26,31^. The native Cx46/50 system, prepared without exogenous sterols and supported by matched physiological reference states, therefore provides a complementary context for distinguishing inhibitor-associated changes from potential effects of the biochemical environment.

### Similarities to physiological NT/TM2 gating

The 2-APB-induced conformational ensemble resembles those observed under low pH and Ca^2+^-gating conditions, suggesting that pharmacological and physiological stimuli converge on a shared NT/TM2 conformational landscape^22,30^. Consistent with low pH, 2-APB treatment produced mixtures of destabilized-open and gated subunits within individual channels. The distributions and spatial arrangements of these conformations were consistent with independent subunit occupancy, producing asymmetric NT-gating ensembles rather than a concerted channel-wide transition (**Supplementary Fig. 6**). This interpretation is also consistent with the near-unity Hill coefficients reported for Cx46 and Cx50 inhibition by 2-APB (1.2 and 0.9, respectively) and functional evidence that individual subunits can initiate channel gating^35,54^. Across all three conditions, displacement of the NT toward the pore is coupled to movement of the cytoplasmic portion of TM2 around the conserved P88 kink, a region that has also been previously implicated in CO_2_ induced gating and voltage sensing mechanism of Cx26 and Cx32^28,57,58^.

The relationship between NT displacement and lipid occupancy also shows parallels across these gating conditions. Whereas low-pH gating appears intrinsically coupled to pore-lipid occupancy, both Ca^2+^ and 2-APB can displace the NT from its open-state position in amphipol without added phospholipid. In nanodiscs, however, both treatments produce ordered gated conformations accompanied by pore-bound densities resembling those assigned as lipids in the low-pH-gated structure. The positions of these densities overlap, although their packing arrangements differ, suggesting that lipids can adopt multiple configurations capable of stabilizing an inward-displaced NT rather than forming a single, conformationally defined “gasket”. The recurring interstitial lipid densities in the low pH and 2-APB-gated states, together with narrowing of the corresponding fenestration in the CBX-flex state, further raise the possibility that this region provides a route for lipid exchange between the bilayer and pore. However, whether this site serves as the point of lipid translocation remains to be defined. Collectively, these observations support convergence on a shared downstream NT/TM2 architecture while allowing pH, Ca^2+^, inhibitors, and lipids to make distinct contributions to its initiation and stabilization.

### Structural basis of broad inhibitor activity and implications for selectivity

The combination of a shared, shallow hydrophobic core and sequence-variable margins of the NT-docking pocket helps explain both broad inhibitor activity as well as potential opportunities for selectivity. Conserved core interactions can accommodate chemically diverse ligands, whereas variable residues may influence binding or couple pocket occupancy to different NT responses. A39 in Cx46 and T39 in Cx50 provide one candidate source of their differential sensitivity to 2-APB, while substitution of the equivalent Cx32 residue with tryptophan produced functional channels with markedly reduced sensitivity to both mefloquine and 2-APB^45^. Selective modulation may therefore require interactions with variable pocket margins or preferential stabilization of isoform-specific conformational states rather than engagement of the conserved hydrophobic core alone. Notably, while conservation of this pocket helps explain broad activity among connexin isoforms, it does not account for the off-target actions of 2-APB and CBX on unrelated protein families^41,59–62^.

### Limitations and Outlook

Several limitations constrain mechanistic interpretation of these structures. The ligand-associated densities do not define unique binding poses. CBX can be accommodated in multiple orientations, whereas the 2-APB-associated density may contain 2-APB, its hydrolysis product diphenylborinic acid, phospholipid, or heterogeneous combinations of these species. Although the amphipol preparation contained no added phospholipid, residual native lipids co-purified with the channels cannot be excluded. The high inhibitor concentrations used to promote structural occupancy for structural analysis may also favor low-affinity poses, heterogeneous binding, or effects that are less prominent near functional IC_50_ values. Furthermore, the resolved conformational ensembles cannot yet be directly assigned to defined conductance states. Mechanistic validation by mutagenesis is intrinsically complicated because the inhibitor-sensitive pocket is also the native NT-docking site; as such, mutations designed to disrupt inhibitor binding could independently alter basal conductance, open-state stability, or gating. Addressing these uncertainties will require additional connexin–inhibitor structures, direct functional measurements, and systematic evaluation of how the membrane environment contributes to pore occlusion, inhibitor occupancy and conformational responses.

## Acknowledgements

We thank Dr. Claudia López and the staff of the OHSU Multiscale Microscopy Core for training and instrumentation access. We also acknowledge computational support from the OHSU Advanced Computing Center (NIH S10OD034224) and cryo-EM instrumentation and user support from the Pacific Northwest Cryo-EM Center (R24GM154185). This work was supported by NIH R35GM124779 (S.L.R.).

## Author Contributions

J.M.J. designed and performed the 2-APB and CBX binding experiments. J.M.J. and J.B.M. collected the cryo-EM data and processed the 2-APB nanodisc dataset. J.M.J. processed the CBX and 2-APB amphipol datasets and built all atomic models. All authors contributed to data analysis and interpretation. J.M.J. and S.L.R. prepared the initial manuscript, and all authors reviewed and edited the final manuscript.

## Methods

### Membrane scaffold protein expression and purification

His-tagged membrane scaffold protein MSP1E1 was expressed in *E. coli* from a plasmid obtained from Addgene^63,64^, and purified as previously described^21,22^. Briefly, cell pellets were resuspended in 40 mM Tris (pH 7.4), 150 mM NaCl, 1% Triton X-100 and 1 mM PMSF and stored at −80 °C. MSP1E1 was batch-bound from clarified lysate to Ni-NTA resin for 1 h at 4 °C with gentle rotation. After washing, MSP1E1 was eluted with 40 mM Tris (pH 8.0), 300 mM NaCl and 500 mM imidazole and further purified by size-exclusion chromatography (SEC) using an ENrich SEC 70 column on an NGC chromatography system (Bio-Rad) at 4 °C. The column was equilibrated in 20 mM HEPES (pH 7.4), and 150 mM NaCl. Fractions containing MSP1E1 were identified by silver-stained SDS-PAGE, pooled and concentrated to 300 μM, as determined by UV_280_ absorbance at 280 nm, using a 5 kDa molecular-weight-cutoff (m.w.c.o.) spin concentrator (Vivaspin 6, Sartorius). Aliquots were flash-frozen in liquid nitrogen and stored at −80 °C.

### Purification of native Cx46/50 from lens tissue

Fresh *ovine* eyes were obtained from Nebraska Scientific (Omaha, NE). Lenses were surgically extracted and stored at −80 °C until use. Plasma membranes from the lens cores, enriched in C-terminally truncated Cx46/50^48–50^, were isolated, stripped and solubilized as previously described^20–22,24,30^. Membranes were solubilized in 1% (wt vol^-1^) *n*-decyl-β-D-maltopyranoside (DM) and native Cx46/50 was purified by anion-exchange chromatography using a UNO Q column (Bio-Rad) and eluted with 10 mM Tris, pH 8.0, 350 mM NaCl, 2 mM EDTA, 2 mM EGTA and 0.03% DM. Fractions containing detergent-solubilized Cx46/50 were identified by silver-stained SDS-PAGE, pooled and concentrated to 270 μM (UV_280_) using a 50 kDa m.w.c.o spin concentrator (Vivaspin 6, Sartorius).

### Reconstitution of Cx46/50 into nanodiscs

Freshly purified Cx46/50 was reconstituted into MSP1E1 nanodiscs containing dimyristoylphosphatidylcholine (DMPC) to obtain open-state channels, as described previously^22^. DMPC dissolved in chloroform was dried overnight under vacuum to form a thin film. The lipid film was resuspended to 35 mM in solubilization buffer containing 20 mM HEPES (pH 7.4), 150 mM NaCl, 2 mM EDTA, 2 mM EGTA and 5% DM and sonicated for 40 min. Cx46/50 was mixed with solubilized DMPC at a molar ratio of 0.6:90 (final DMPC concentration, 5 mM) and rotated gently for 1 h at 25 °C. MSP1E1 was then added to obtain a final molar ratio of 0.6:1:90 (Cx46/50:MSP1E1:DMPC), followed by gentle rotation for 15 min at 25 °C.

Detergent was removed overnight by incubation with Bio-Beads SM-2 (Bio-Rad) at a 30:1 bead:DM mass ratio with gentle rotation at 25 °C. The Bio-Beads were removed by centrifugation and replaced with fresh Bio-Beads at the same ratio for an additional 1 h. Insoluble material was removed by centrifugation at 16,000 × *g* for 30 min at 4 °C, and the supernatant was passed through a 0.22 μm filter. The sample was then separated by SEC on a Superose 6 Increase 10/300 GL column (Cytiva) equilibrated in 20 mM HEPES (pH 7.4), and 150 mM NaCl. Fractions containing nanodisc-embedded Cx46/50 were identified by silver-stained SDS-PAGE, pooled and concentrated to ∼4.0 mg mL^-1^ (UV_280_) using a 50 kDa m.w.c.o spin concentrator (Vivaspin 6, Sartorius).

### Reconstitution of Cx46/50 into amphipol

Freshly purified Cx46/50 was exchanged from DM into amphipol A8-35 (Anatrace) to obtain open-state channels, as previously described^20^, with minor modifications. Detergent-solubilized Cx46/50 was mixed with amphipol at a 3:1 amphipol:protein mass ratio and rotated gently overnight at 4 °C. Detergent was removed by incubation with Bio-Beads SM-2 (Bio-Rad) at a 30:1 bead:DM mass ratio for 1 h at 4 °C with gentle rotation. Insoluble material was removed by centrifugation at 16,000 × *g* for 30 min at 4 °C, and the supernatant was passed through a 0.22 μm filter. The sample was separated by SEC on a Superose 6 Increase 10/300 GL column (Cytiva) equilibrated in 20 mM HEPES (pH 7.4), and 150 mM NaCl. Peak fractions containing amphipol-reconstituted Cx46/50 were confirmed by SDS-PAGE, pooled and concentrated to ∼3.0 mg mL^-1^ (UV_280_) using a 50 kDa m.w.c.o spin concentrator (Vivaspin 6, Sartorius).

### Small molecule binding conditions

Freshly prepared Cx46/50 samples were diluted to the concentrations used for negative-stain EM or cryo-EM and treated with either 300 μM 2-APB in 0.5% DMSO (vol vol^-1^) or 5 mM CBX. Samples were incubated on ice for 1 h before application to EM grids. The 300 μM 2-APB concentration is ∼10- and ∼100-fold above the reported IC_50_ values for Cx46 and Cx50, respectively^35^. Because IC_50_ values for CBX have not been reported for Cx46 or Cx50, samples were treated with 5 mM CBX, corresponding to ∼10- to ∼100-fold above published IC_50_ values for other connexin isoforms^36,37,51^.

### Negative-stain electron microscopy

Cx46/50 samples treated with either 300 μM 2-APB or 5 mM CBX were prepared for negative-stain EM as described previously^20–22,24,30^. A 3 μL aliquot of sample at 0.03 mg mL^-1^ was applied to a glow-discharged, continuous carbon-coated EM grid (G400, Ted Pella), blotted with filter paper and washed three times with SEC buffer and then stained with freshly prepared 0.75% (wt vol^-1^) uranyl formate (SPI-Chem). Specimens were imaged at the OHSU Multiscale Microscopy Core using a 120 kV iCorr TEM (Thermo Fisher Scientific) operated at a nominal magnification of 49,000× and equipped with a 2k × 2k Eagle 2K TEM CCD camera (Thermo Fisher Scientific). Images were acquired at a calibrated specimen pixel size of 4.37 Å pixel^-1^ and a nominal defocus of −1.5 μm.

### Cryo-EM specimen preparation and data collection

Cryo-EM samples comprised Cx46/50 in amphipol treated with 300 μM 2-APB or Cx46/50 in nanodiscs treated with either 300 μM 2-APB or 5 mM CBX. For each condition, 3 μL of freshly prepared samples containing Cx46/50 at 1.6–2.0 mg mL^-1^ was applied to glow-discharged Quantifoil holey carbon grids (R1.2/1.3 or R2/1). Grids were prepared using a Vitrobot Mark IV (Thermo Fisher Scientific) at 20 °C and 100% relative humidity, blotted for 4.0–5.5 s, plunge-frozen in liquid ethane and stored under liquid nitrogen. Grids were screened at the Pacific Northwest Cryo-EM Center (PNCC) using a Talos Arctica transmission electron microscope (Thermo Fisher Scientific) operated at 200 kV and equipped with a Gatan K3 detector. Screening data were acquired using SerialEM^65^.

All high-resolution datasets were collected at the PNCC using a Titan Krios G4 TEM (Thermo Fisher Scientific) operated at 300 kV. For 2-APB-treated Cx46/50 in nanodiscs, movies were acquired using a K3 direct electron detector (Gatan) at a nominal magnification of 29,000× in super-resolution mode, corresponding to a calibrated physical pixel size of 0.802 Å pixel^-1^, using a 20 eV energy-filter slit. Automated acquisition with SerialEM yielded 13,202 movies collected with 53 frames, a total dose of 40 e^-^Å^-2^ and a nominal defocus range of −0.5 to −1.8 μm.

For 2-APB-treated Cx46/50 in amphipol, movies were acquired using a Falcon 4i direct electron detector (Thermo Fisher Scientific) at a nominal magnification of 105,000× in super-resolution EER mode, corresponding to a calibrated physical pixel size of 1.176 Å pixel^-1^, using a 10 eV energy-filter slit. Automated acquisition with EPU^66^ (Thermo Fisher Scientific) yielded 7,311 movies with a total dose of 40 e^-^Å^-2^ and a nominal defocus range of −0.5 to −1.8 μm.

For CBX-treated Cx46/50 in nanodiscs, movies were acquired using a Falcon 4i direct electron detector at a nominal magnification of 96,000× in EER mode, corresponding to a calibrated physical pixel size of 0.814 Å pixel^-1^, using 10 eV energy-filter slit. Automated acquisition with EPU yielded 7,748 movies with a total dose of 50 e^-^Å^-2^ and a nominal defocus range of −0.5 to −2.5 μm.

### Image processing of 2-APB-treated Cx46/50 in nanodisc

The 2-APB dataset was processed using CryoSPARC (v4.7.1)^67,68^ (**Supplementary Fig. 2**). Movies were dose-weighted and corrected for beam-induced motion using Patch Motion Correction. Micrographs collected in super-resolution mode were 2x binned during motion correction to the calibrated physical pixel size of 0.802 Å pixel^-1^, hereafter referred to as unbinned sampling. Contrast transfer function (CTF) parameters were estimated using Patch CTF.

An initial 3D template was generated from ∼500,000 blob-picked particles through three rounds of 2D classification followed by *ab initio* reconstruction. The map was subjected to non-uniform (NU) refinement with D6 symmetry (1.60 Å voxel size) and low-pass filtered for template-based particle picking. Template picker assigned ∼7.2 million initial particle picks that were curated through two rounds of multiclass *ab initio* reconstruction. Classes displaying gap junction features were combined (∼1.3 million particles), NU-refined and subjected to three rounds of heterogeneous refinement with progressive unbinning. This yielded two high-resolution particle stacks distinguished by their NT conformations (open-like and gated-like), which were processed independently.

#### (i) 2-APB-treated destabilized-open state

The open-like particle stack (230,094 particles) was subjected to unbinned NU-refinement with D6 symmetry, yielding a 2.2 Å reconstruction. Because the NT density remained discontinuous, particles were binned to 1.70 Å pixel^-1^, expanded according to D6 symmetry (∼2.8 million symmetry-expanded particles) and subjected to two rounds of 3D variability analysis (3DVA), using a custom mask generated in ChimeraX^69^ to exclude nanodisc density.

Each principal component (PC) was divided into nine frames with a rolling window of zero. The three central frames (frames 3–5), which contained the most coherent open-position NT density, were selected from each PC. For each PC, the selected particles were combined, symmetry expansion was undone, and the particles were returned to unbinned sampling for NU-refinement with D6 symmetry. The three PC-selected particle stacks were then combined and deduplicated, yielding 165,916 particles. Four-class 3D classification with hard assignments using the 3DVA mask identified one class of 49,198 particles with a well-resolved destabilized-open NT conformation. This class was refined to 2.4 Å resolution by gold-standard FSC and used for atomic modeling (**Supplementary Fig. 3; Supplementary Table 1**).

#### (ii) 2-APB-treated gated state

The gated-like particle stack (328,379 particles) was subjected to unbinned NU-refinement with D6 symmetry, yielding a 2.1 Å reconstruction. The NT displayed gated-like features at low contour levels but retained open-like features at higher contour levels, indicating residual heterogeneity. Particles were therefore binned to 1.70 Å pixel^-1^, NU-refined with D6 symmetry, symmetry expanded (∼3.9 million particles) and subjected to two rounds of 3DVA using the mask described above. The first round removed particles at PC extremes that obscured NT variability. The second identified a PC spanning fully open and fully gated NT conformations. Particles at the gated endpoint (frame 0) were selected, symmetry expansion was undone and the resulting 25,706 particles were returned to unbinned sampling for NU refinement with D6 symmetry.

To further enrich the gated-state population, particles were rebinned to 1.70 Å pixel^-1^, NU-refined with D6 symmetry and symmetry expanded (∼308,000 particles). Signal outside a single connexin subunit was subtracted using a custom ChimeraX mask generated from merged open-state (PDB 7JKC) and gated-state (PDB 9Z9X) Cx46 models. Three-class 3D classification with hard assignments identified one class with resolved gated NT density, comprising ∼33% of the particles. Signal subtraction and symmetry expansion were undone, yielding 8,978 particles for final unbinned NU refinement with D6 symmetry. The resulting 2-APB-gated reconstruction reached 2.8 Å resolution by gold-standard FSC and was used for atomic modeling (**Supplementary Fig. 3; Supplementary Table 1**).

#### (iii) Subunit analysis of the 2-APB consensus particle stack

The consensus particle stack (721,863 particles) was binned 1.70 Å pixel^-1^ and symmetry expanded according to D6 symmetry. Signal outside an individual connexin subunit was subtracted using a custom mask generated in ChimeraX from superimposed open-state (PDB 7JKC) and gated-state (PDB 9Z9X) Cx46 subunit models. The isolated subunits were then subjected to 3DVA, which identified a PC spanning open and gated NT conformations. Particles associated with the open-like and gated-like regions of this component were assigned to the corresponding states by visual inspection, yielding ∼60% open and 40% gated subunits. Particle stacks representing each conformation were combined separately and refined. Subunit-state assignments were then mapped back to their parent gap junction particles using the previously described *Metadata Extraction, Gap Junction Splitter, Subunit State Frequency* and *Hemichannel Arrangement* scripts^22^. Observed arrangement frequencies were compared with those expected under random subunit occupancy, and the resulting frequency and arrangement distributions were plotted for visualization (**Supplementary Fig. 6**).

### Image processing for 2-APB-treated Cx46/50 in amphipol

The amphipol dataset was processed using CryoSPARC (v4.7.1) (**Supplementary Fig. 4**). EER movies were imported as 40 fractions, up-sampled two-fold, and corrected for beam-induced motion using Patch Motion Correction with an F-crop factor of 1, yielding a super-resolution pixel size of 0.588 Å pixel^-1^. CTF parameters were estimated using Patch CTF.

An initial 3D template was generated from ∼107,000 blob-picked particles through two rounds of 2D classification followed by *ab initio* reconstruction. The particles were subjected to NU-refinement with D6 symmetry, and the resulting map was low-pass filtered for 3D template-based particle picking. Template picker yielded ∼6 million initial particle picks, which were cleaned through two rounds of 2D classification and multiclass *ab initio* reconstruction. Classes displaying gap junction features were combined, yielding ∼283,000 particles. NU refinement with D6 symmetry produced a 2.6 Å consensus reconstruction (gold-standard FSC).

The consensus particles were binned to 1.73 Å pixel^-1^ and subjected to three iterative rounds of 3DVA using a custom mask generated in ChimeraX that excluded the amphipol micelle. In each round, every PC was divided into nine frames with a rolling window of zero. The central frame (frame 4), which contained the most consistently resolved gated-state NT density, was selected from each PC. The selected particles were returned to unbinned sampling, NU-refined and combined across PCs before the next round. After the third round, duplicate particles were removed, yielding 29,546 particles for final NU-refinement with D6 symmetry. The resulting 2-APB-gated reconstruction reached 3.2 Å resolution (gold-standard FSC) and was used for atomic modeling (**Supplementary Fig. 5; Supplementary Table 1**).

### Image processing of CBX-treated Cx46/50 in nanodisc

The CBX dataset was processed using CryoSPARC (v4.7.1) (**Supplementary Fig. 7**). EER movies were imported as 50 fractions with an up-sampling factor of 1. Movies were dose-weighted and corrected for beam-induced motion using Patch Motion Correction, yielding a pixel size of 0.814 Å pixel^-1^. CTF parameters were estimated using Patch CTF.

An initial 3D template was generated from ∼1.1 million blob-picked particles through three rounds of 2D classification followed by *ab initio* reconstruction. The map was subjected to NU-refinement with D6 symmetry (1.60 Å voxel size) and low-pass filtered for template-based particle picking. Template picker assigned ∼4.5 million initial particle picks, which were cleaned through three rounds of multiclass *ab initio* reconstruction. Classes displaying gap junction features were combined (∼290,000 particles), returned to unbinned sampling and subjected to NU-refinement followed by six-class heterogeneous refinement. Four classes with high-resolution features were combined (∼243,000 particles) and subjected to NU-refinement with D6 symmetry, yielding a 2.4 Å consensus reconstruction.

Conformational heterogeneity within the consensus particle stack was examined using 3DVA and heterogeneous refinement. For 3DVA, particles were binned to 1.90 Å pixel^-1^, expanded according to D6 symmetry and analyzed using a custom mask generated in ChimeraX that excluded nanodisc density. The second PC suggested NT variability consistent with pairing between neighboring subunits. However, reconstructions along this component did not resolve clear NT conformations when refined with D6, D3, D2, C6 or C1 symmetry. Ultimately, four-class heterogeneous refinement of the unbinned consensus particles identified two nearly identical high-resolution classes. Combining these particle stacks did not improve reconstruction quality or NT resolvability. A single class containing 87,277 particles was therefore selected for final NU-refinement with D6 symmetry. The resulting CBX-flex reconstruction reached 2.8 Å resolution (gold-standard FSC) and was used for atomic modeling (**Supplementary Fig. 8; Supplementary Table 1**).

### Atomic modeling and validation

Previously published open-state models of Cx46 (PDB 7JKC) and Cx50 (PDB 7JJP) were used as starting models for atomic modeling in Coot^70^. Raw and locally filtered cryo-EM maps were used during model building, and final models were validated against the locally filtered maps. For the 2-APB-gated and CBX-flex reconstructions, NT residues preceding position 20 were removed from the starting models and rebuilt where supported by density.

Models were iteratively adjusted and refined using Coot, ISOLDE^71^ and Phenix^72^ until convergence of statistics as determined by MolProbity^73,74^ (**Supplementary Table 1**). All final models omit the intracellular loop (ICL; residues 105–138 in Cx46 and 105–151 in Cx50) and C-terminal residues after position 223 in Cx46 or 235 in Cx50 due to lack of resolvability of these regions in the density maps. In the 2-APB-gated models, NT residues 2–5 were omitted and the side chains of residues 6–17 were truncated to Cβ because of limited local resolution. In the CBX-flex models, NT residues 2–17 were omitted because no corresponding density was resolved.

DMPC molecules were fitted into lipid-like densities in Coot, and atoms extending beyond the supported density were removed. Candidate poses of 2-APB and CBX^75,76^ were placed into their corresponding putative densities in Coot for visualization. Candidate CBX poses were additionally evaluated using AutoDock Vina docking-score calculations, as described below. These ligand assignments remain tentative and are not included in the final deposited models.

Coulombic surface potentials were calculated, analyzed and visualized in ChimeraX. Sequence alignments were generated using Clustal-w^77^ and visualized in Jalview^78^.

### AutoDock scoring

AutoDock Vina (v1.2.0)^55^ was used to estimate docking energies for CBX in the Cx46 and Cx50 CBX-flex models. Three candidate CBX conformations were fitted to features of the CBX-compatible cryo-EM density in Coot and evaluated separately. For each analysis, the corresponding CBX-flex model was reduced to a dimer of adjacent connexin subunits, lipids were removed, and the candidate CBX conformation was saved as a separate PDB file. Protein and ligand input files were prepared using MGLTools (v1.5.7)^79,80^.

The docking search space was restricted to the TM1/TM2 groove encompassing the CBX-compatible density using a grid box of 20 × 30 × 32 Å centered at x = 169, y = 147 and z = 200 Å, with a grid spacing of 1 Å. The internal geometry and rotatable bonds of each density-fitted CBX conformation were held fixed during docking. For each candidate conformation, the top-ranked Vina docking-energy estimate was reported.

### AI-assisted technologies

During the development of this manuscript ChatGPT (OpenAI) was used to revise portions of the text to improve clarity. The authors reviewed and edited any content that was revised with this tool and take full responsibility for the content of this publication.

## Data availability

Cryo-EM density maps have been deposited to the Electron Microscopy Data Bank (EMDB) under accession numbers: EMD-XXXXX (2-APB gated state, nanodiscs), EMD-XXXXX (2-APB destabilized-open state connexin-46/50, nanodiscs), EMD-XXXXX (2-APB gated state connexin-46/50, amphipol) and EMD-XXXXX (CBX flex state connexin-46/50, nanodiscs). Coordinates for atomic models have been deposited to Protein Data Bank (PDB) under accession numbers: XXXX (2-APB gated state connexin-46, nanodiscs), XXXX (2-APB gated state connexin-50, nanodiscs), XXXX (2-APB destabilized-open state connexin-46, nanodiscs), XXXX (2-APB destabilized-open state connexin-50, nanodiscs), XXXX (2-APB gated state connexin-46, amphipol), XXXX (2-APB gated state connexin-50, amphipol), XXXX (CBX flex state connexin-46, nanodisc) and XXXX (CBX flex state connexin-50, nanodisc). The original raw multi-frame micrographs have been deposited to EMPIAR under accession numbers: EMPIAR-XXXXX (2-APB treated connexin-46/50, nanodiscs), EMPIAR-XXXXX (2-APB treated connexin-46/50, amphipol) and EMPIAR-XXXXX (CBX treated connexin-46/50, nanodiscs). Previously published models used for modeling and comparative analysis can be found at the PDB: 7JKC and 7JJP. Previously published maps used for comparative analysis can be found at: Cx46/50 (EMD-73962), Cx32 (EMD-18446), Cx36 (EMD-38322, EMD-38327, EMD-18540 and EMD-38320).

## Supplementary Figures and Tables

**Supplementary Figure 1:**
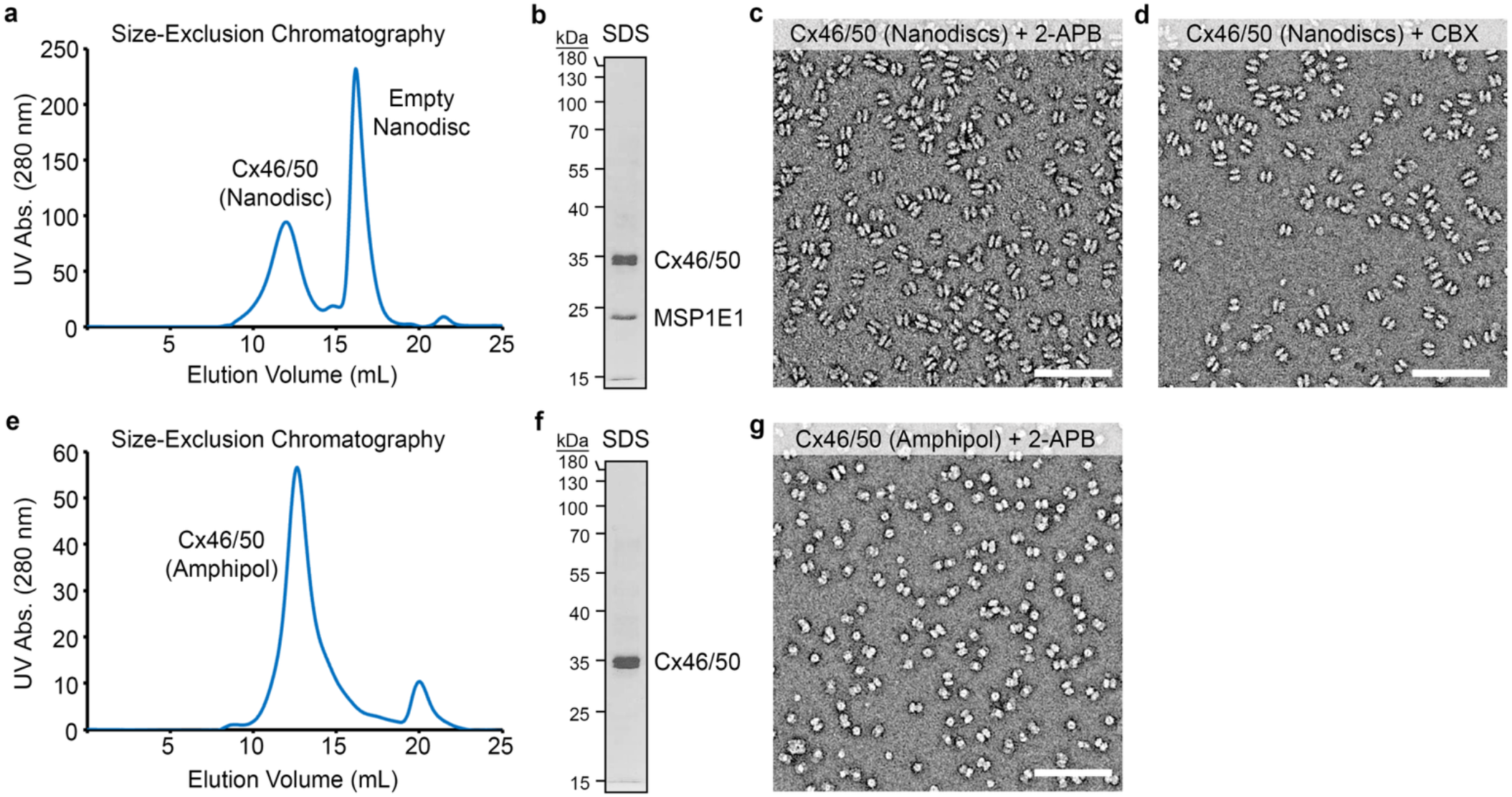
Purification and negative-stain EM of inhibitor-treated Cx46/50 in nanodiscs and amphipol. **a**, Size-exclusion chromatography (SEC) monitored by absorbance at 280 nm, with peaks containing native Cx46/50 gap junction channels reconstituted in MSP1E1–DMPC nanodiscs and empty nanodiscs are indicated. **b,** Silver-stained SDS–PAGE of the Cx46/50-nanodisc peak. MSP1E1 migrates at ∼24 kDa (predicted molecular weight, ∼27.5 kDa), whereas native lens Cx46 and Cx50 isolated from core tissue co-migrate at ∼38 kDa. **c,d,** Negative-stain electron micrographs of nanodisc-reconstituted Cx46/50 following treatment with (*c*) 300 µM 2-APB or (*d*) 5 mM CBX. **e,** SEC of native Cx46/50 in amphipol monitored at 280 nm, with the Cx46/50-containing peak indicated. **f,** Silver-stained SDS–PAGE of the primary peak, showing Cx46/50 at ∼38 kDa. **g,** Negative-stain electron micrograph of amphipol-reconstituted Cx46/50 following treatment with 300 µM 2-APB. Scale bars, 100 nm (panels *c,d,g*). Molecular weight (MW) markers for panels *(b)* and *(f)* are indicated in kDa.

**Supplementary Figure 2:**
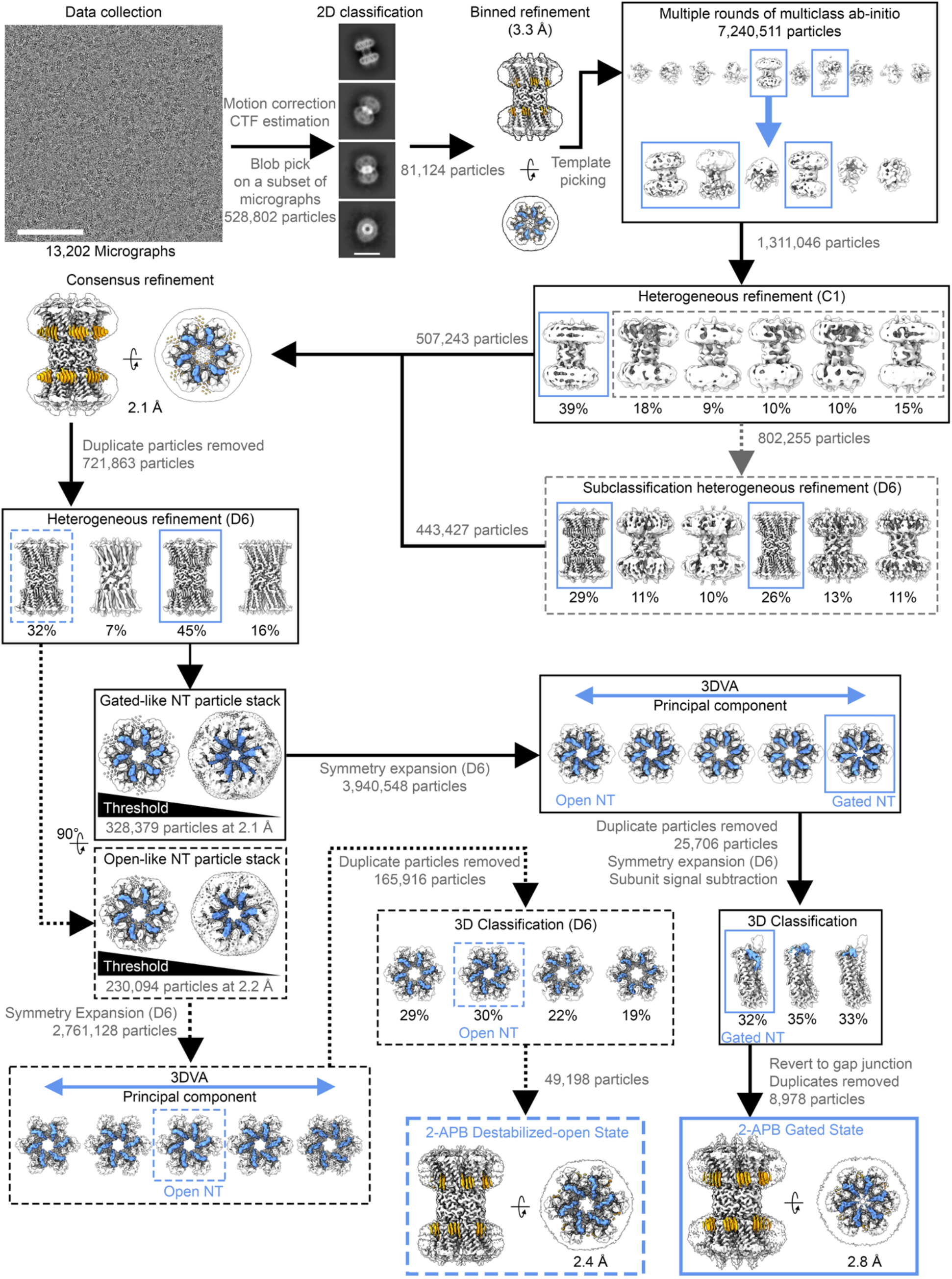
Image processing workflow for 2-APB-treated Cx46/50 in nanodiscs. Image processing workflow for Cx46/50 in DMPC nanodiscs treated with 300 µM 2-APB performed in CryoSPARC. A representative cryo-EM micrograph and selected 2D class averages are shown (scale bars, 100 nm and 100 Å, respectively). Initial blob picking followed by 2D classification and non-uniform refinement produced a 3D template for template-based particle picking. Iterative multiclass *ab initio* reconstruction and heterogeneous refinement performed with and without applied D6 symmetry isolated gap junction particles, yielding a 2.1 Å consensus reconstruction with resolved NT and annular lipid densities (highlighted in blue and orange, respectively). Heterogeneous refinement with D6 symmetry separated gated-like and open-like particle stacks. The gated-like class contained threshold-dependent open- and gated-like NT features, whereas the open-like class retained NT density across thresholds. Both stacks were symmetry expanded and analyzed by 3D variability analysis (3DVA). For the gated-like stack, 3DVA identified a principal component spanning open-like to gated-like NT conformations. Particles at the gated endpoint were selected and refined, then symmetry expanded and signal subtracted to isolate individual connexin subunits for 3D classification. A class with resolved gated NT density was retained containing 8,978 particles, after which signal subtraction and symmetry expansion were undone for final refinement of the 2-APB-gated state to 2.8 Å. For the open-like stack, particles with well-defined open-state NT density identified near the center of the principal components were selected and subjected to 3D classification. A class containing 49,198 particles with resolved NT density was retained for final refinement of the destabilized-open state to 2.4 Å.

**Supplementary Figure 3:**
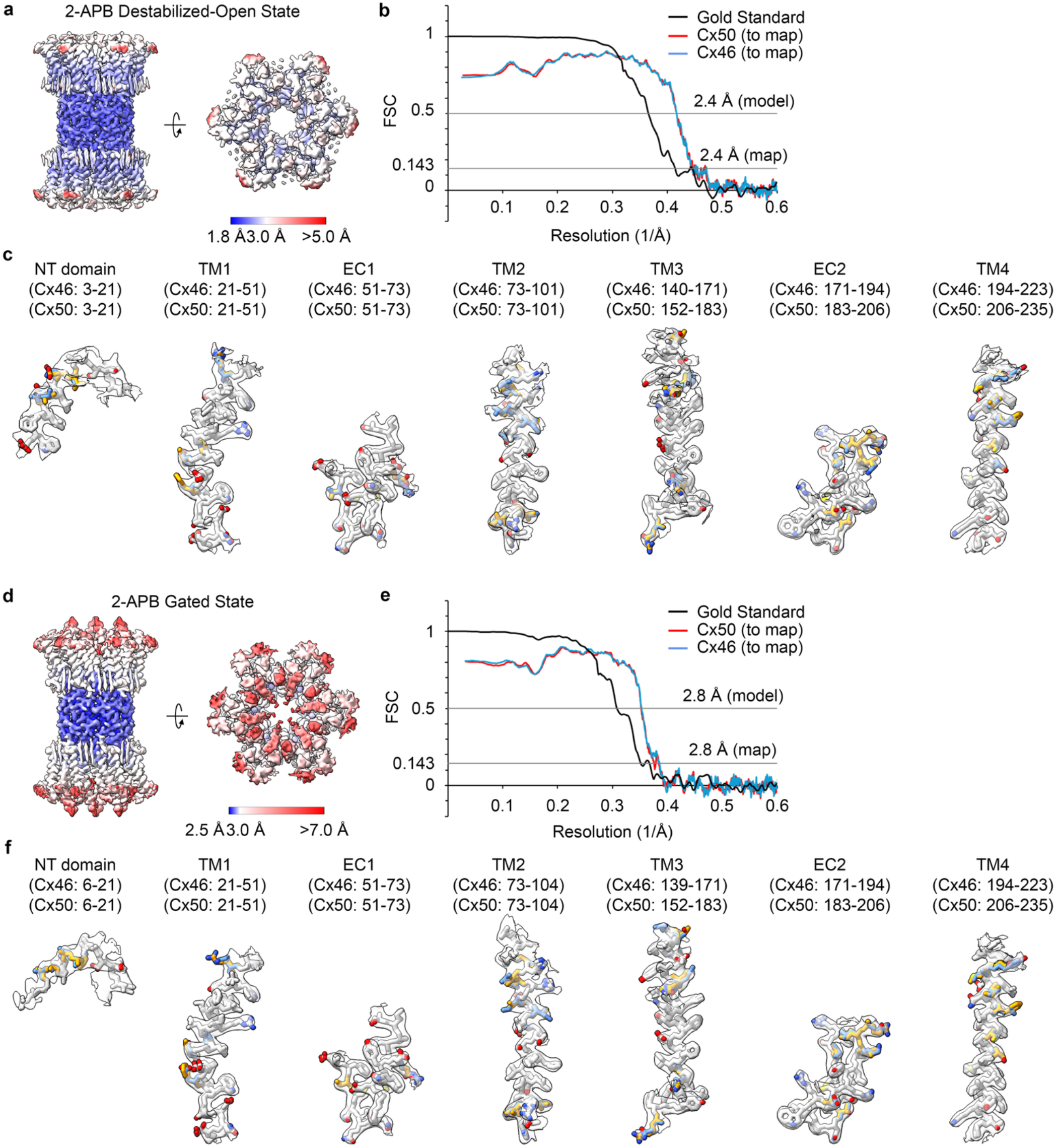
Resolution assessments and atomic-model validation for the 2-APB-gated and destabilized-open states in nanodiscs. **a-c**, Analysis of the destabilized-open state. **d-f,** Analysis of the 2-APB-gated state. **a,d,** Local-resolution estimates of the final cryo-EM reconstructions calculated in CryoSPARC and displayed as colored surfaces (scale shown). **b,e,** Fourier Shell Correlation (FSC) curves for the final cryo-EM reconstructions, with gold-standard FSC (black) indicating global resolutions (0.143 cutoff). Map-to-model FSC curves are shown for Cx46 in blue and Cx50 in red (0.5 cutoff). **c,f,** Segmented cryo-EM densities with fitted Cx46 and Cx50 atomic models for the NT domain, transmembrane helices (TM1–TM4) and extracellular domains (EC1–2). Residues identical between Cx46 and Cx50 are shown in white; isoform-specific residues are shown in blue for Cx46 and orange for Cx50.

**Supplementary Figure 4:**
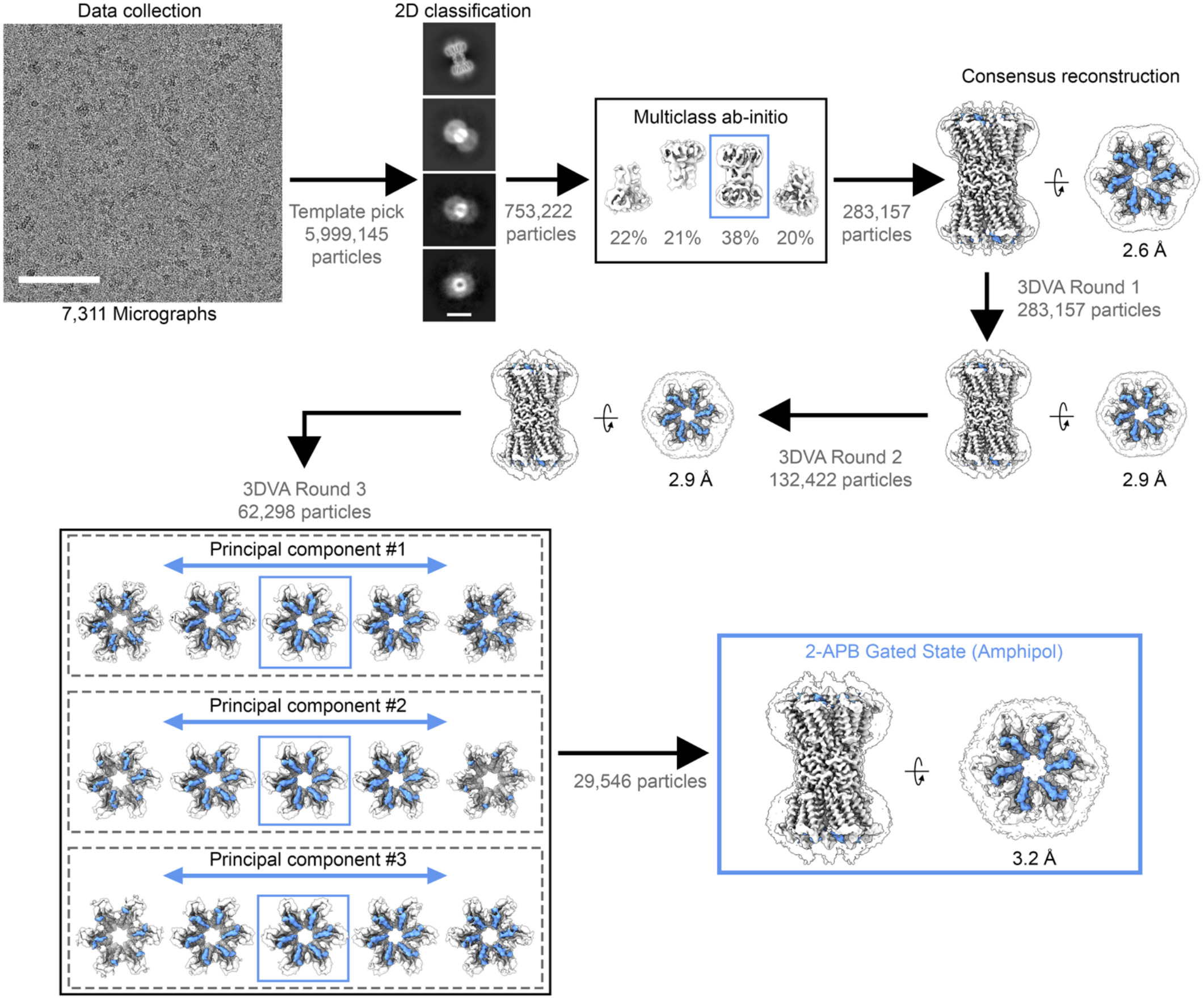
Image processing workflow for 2-APB-treated Cx46/50 in amphipol. Image processing workflow for Cx46/50 in amphipol A8-35 following treatment with 300 µM 2-APB performed in CryoSPARC. A representative cryo-EM micrograph and selected 2D class averages are shown (scale bars, 100 nm and 100 Å, respectively). Initial blob picking, 2D classification and *ab initio* reconstruction generated a template that was refined with D6 symmetry for template-based particle picking (not shown). Template-picked particles were subjected to 2D classification and multiclass *ab initio* reconstruction to isolate gap junction particles. Non-uniform refinement with D6 symmetry yielded a 2.6 Å consensus reconstruction with resolved NT densities (highlighted in blue). The consensus particles were then subjected to three sequential rounds of 3D variability analysis (3DVA). After each round, particles associated with resolved gated NT density near the centers of the principal components were selected, refined, combined and deduplicated before the next round. Following the third round, 29,546 particles were retained for final refinement of the 2-APB-gated state to 3.2 Å.

**Supplementary Figure 5:**
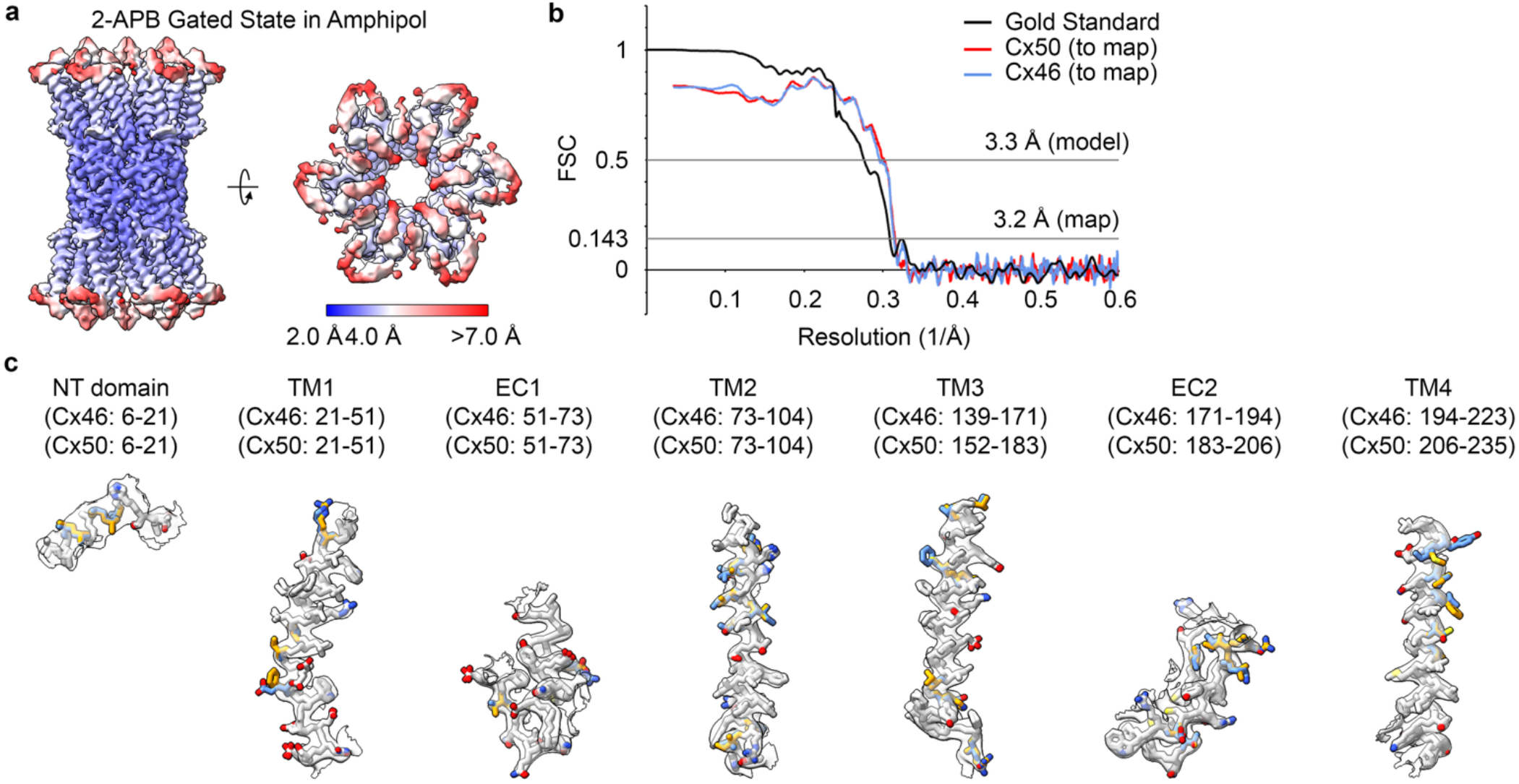
Resolution assessment and atomic model validation for the 2-APB-gated state in amphipol. **a**, Local-resolution estimates of the final cryo-EM reconstruction calculated in CryoSPARC and displayed as colored surfaces (scale shown). **b,** Fourier Shell Correlation (FSC) curves for the final cryo-EM reconstruction, with gold-standard FSC (black) indicating global resolutions (0.143 cutoff). Map-to-model FSC curves are shown for Cx46 in blue and Cx50 in red (0.5 cutoff). **c,** Segmented cryo-EM densities with fitted Cx46 and Cx50 atomic models for the NT domain, transmembrane helices (TM1–TM4) and extracellular domains (EC1–2). Residues identical between Cx46 and Cx50 are shown in white; isoform-specific residues are shown in blue for Cx46 and orange for Cx50.

**Supplementary Figure 6:**
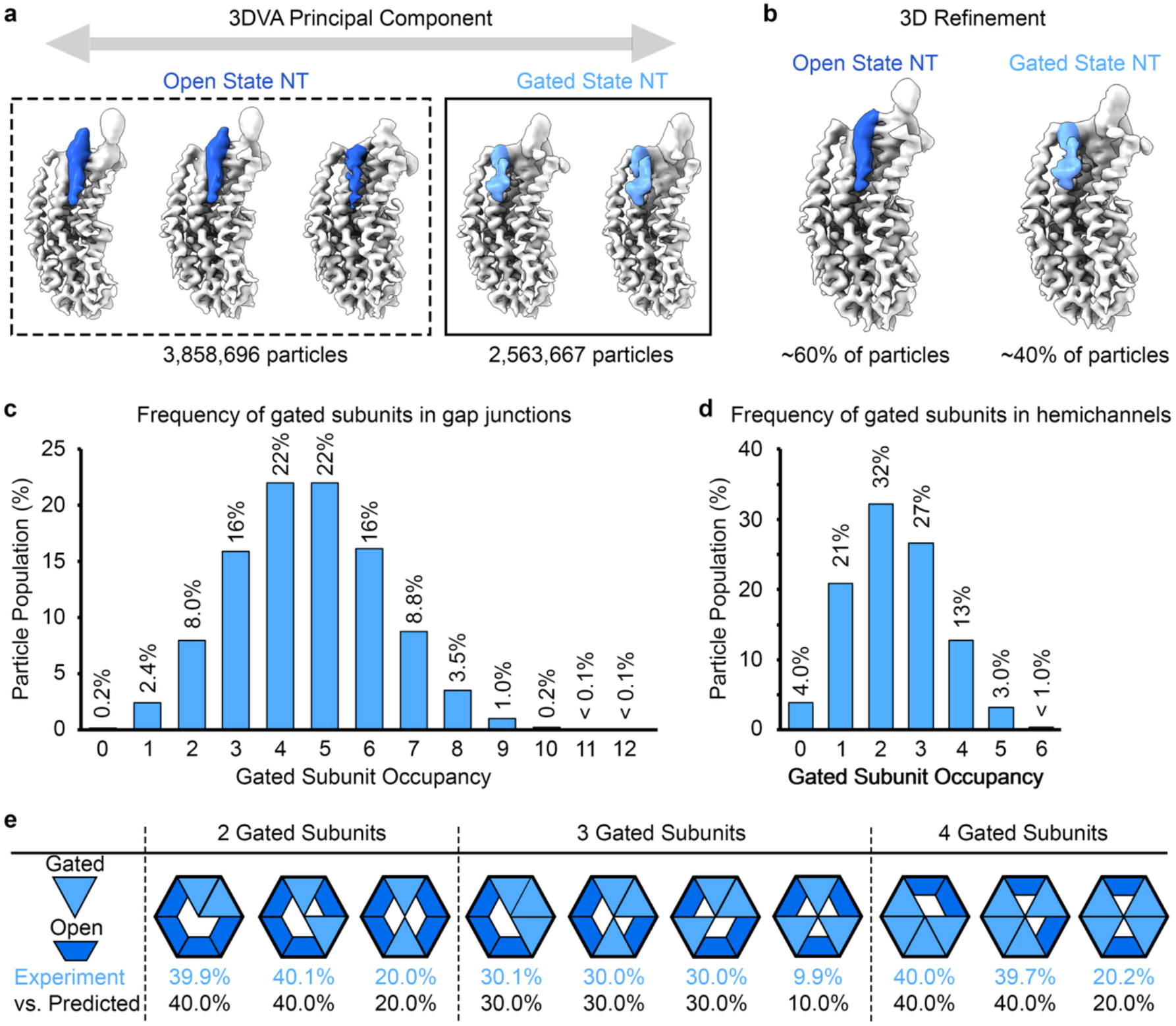
2-APB treatment produces an ensemble of open and gated states. **a**, Consensus particles were expanded according to D6 symmetry, and signal outside an individual connexin subunit was subtracted before 3DVA. The principal component shown spans open and gated NT conformations. **b,** Reconstructions generated by combining particles assigned to the *(left)*, open-like and *(right)*, gated-like regions of the principal component, representing ∼60% and 40% of the assigned subunits, respectively. **c,d,** Frequency distributions of the number of gated subunits within individual *(c)* gap junction channels and *(d)* hemichannels. Gap junctions containing four or five gated subunits were most abundant, each accounting for ∼22% of particles, whereas hemichannels containing two gated subunits were most abundant, accounting for ∼32%. **e,** Distinct spatial arrangements of open (dark blue trapezoids) and gated (light blue triangles) subunits within hemichannels containing two, three or four gated subunits. Observed frequencies are shown in light blue and compared with frequencies predicted under random subunit occupancy in black.

**Supplementary Figure 7:**
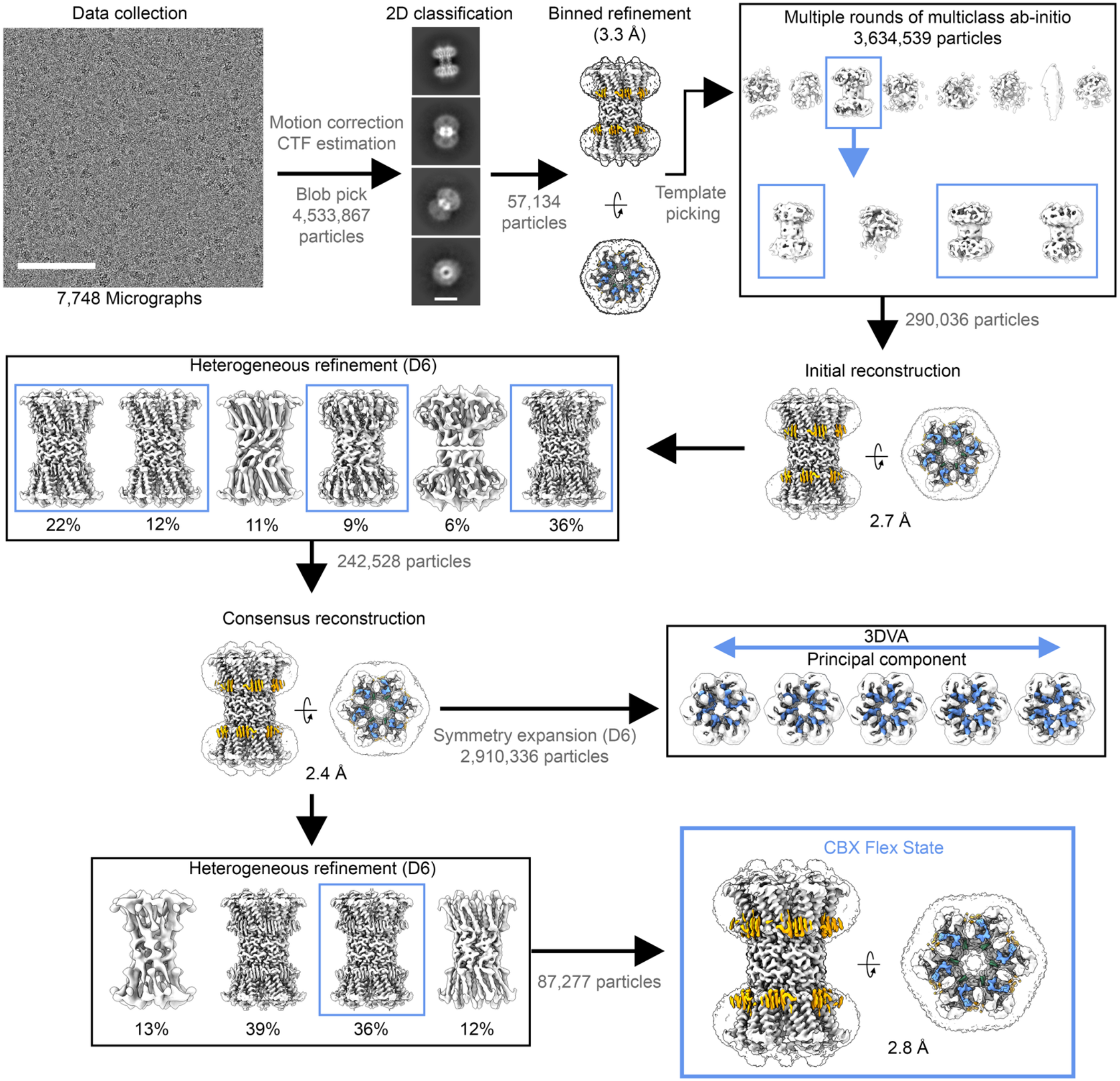
Image processing workflow for CBX-treated Cx46/50 in nanodiscs. Image processing workflow for Cx46/50 in DMPC nanodiscs following treatment with 5 mM CBX performed in CryoSPARC. A representative cryo-EM micrograph and selected 2D class averages are shown (scale bars, 100 nm and 100 Å, respectively). Initial blob picking and 2D classification produced a 3D template for template-based particle picking. Iterative multiclass *ab initio* reconstruction isolated gap junction particles, which were refined and then subjected to heterogeneous refinement with D6 symmetry. Four classes with high-resolution features were combined and subjected to non-uniform refinement with D6 symmetry, yielding a 2.4 Å consensus reconstruction with the hinge-region of the NT and annular lipid densities (highlighted in blue and orange, respectively). To assess conformational heterogeneity, the consensus particles were binned, symmetry expanded and subjected to 3D variability analysis (3DVA). Although variability was observed within the NT region, no discrete ordered NT conformation was resolved upon further refinement. In parallel, heterogeneous refinement of the consensus particles yielded two similar high-resolution classes. One class containing 87,277 particles was ultimately selected for final non-uniform refinement, producing the 2.8 Å CBX-flex state reconstruction.

**Supplementary Figure 8:**
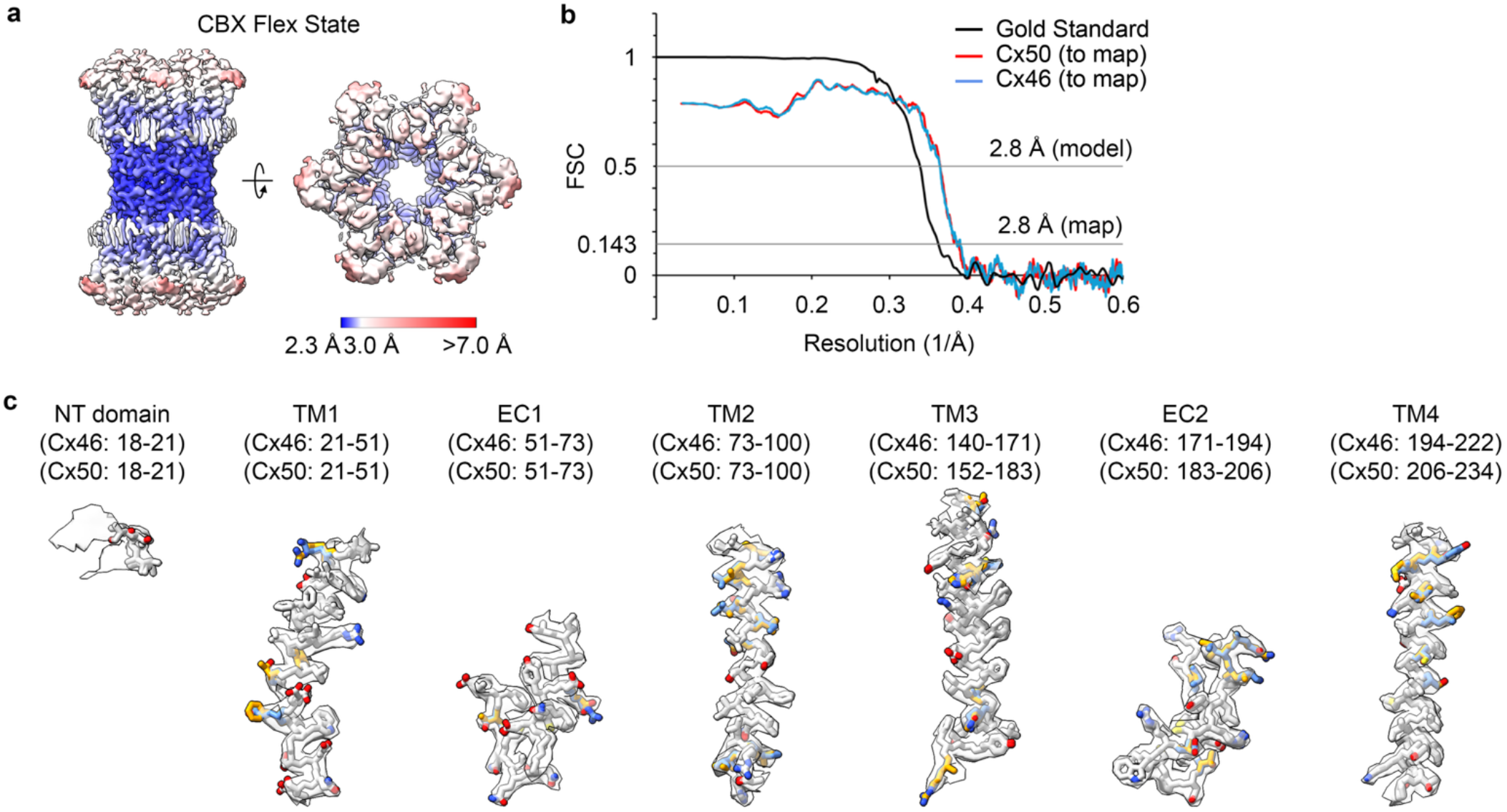
Resolution assessment and atomic model validation for the CBX-flex state in nanodiscs. **a**, Local-resolution estimates of the final cryo-EM reconstruction calculated in CryoSPARC and displayed as colored surfaces (scale shown). **b,** Fourier Shell Correlation (FSC) curves for the final cryo-EM reconstruction, with gold-standard FSC (black) indicating global resolutions (0.143 cutoff). Map-to-model FSC curves are shown for Cx46 in blue and Cx50 in red (0.5 cutoff). **c,** Segmented cryo-EM densities with fitted Cx46 and Cx50 atomic models for the NT domain, transmembrane helices (TM1–TM4) and extracellular domains (EC1–2). Residues identical between Cx46 and Cx50 are shown in white; isoform-specific residues are shown in blue for Cx46 and orange for Cx50.

**Supplementary Figure 9:**
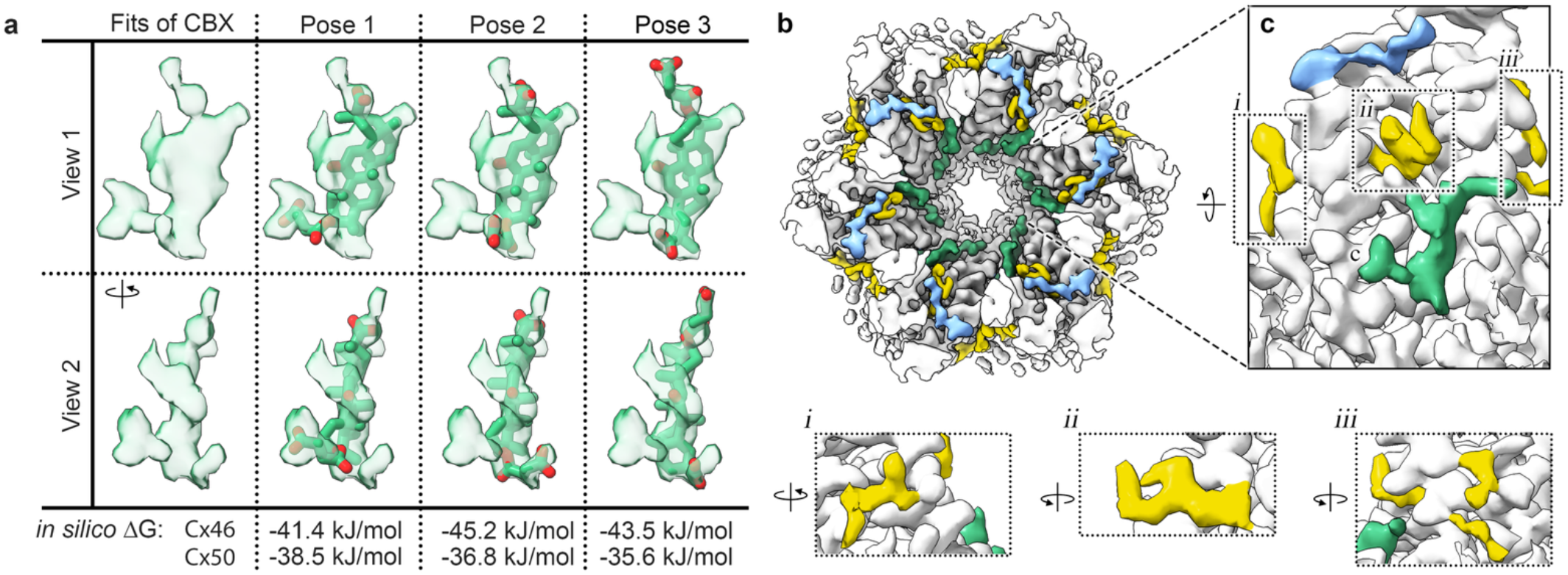
Candidate CBX poses and non-protein densities in the CBX-flex state. **a**, Orthogonal views of the segmented CBX-compatible density shown alone and with three candidate CBX poses. AutoDock Vina docking scores for each pose are indicated. **b,** Cytoplasmic view of the CBX-flex cryo-EM map, with NT-region density shown in blue, CBX-compatible density in green and unassigned densities in yellow. **c,** Rotated close-up of the boxed region in (*b*), showing the unassigned densities relative to the CBX-compatible density. Insets show unassigned densities: *i* near the cytoplasmic leaflet at the TM1/TM4 interface, *ii* within the pore between TM1 and TM2 and positioned between the NT region and CBX-compatible density, and *iii* near the cytoplasmic leaflet at the TM2/TM3 interface.

**Supplementary Figure 10:**
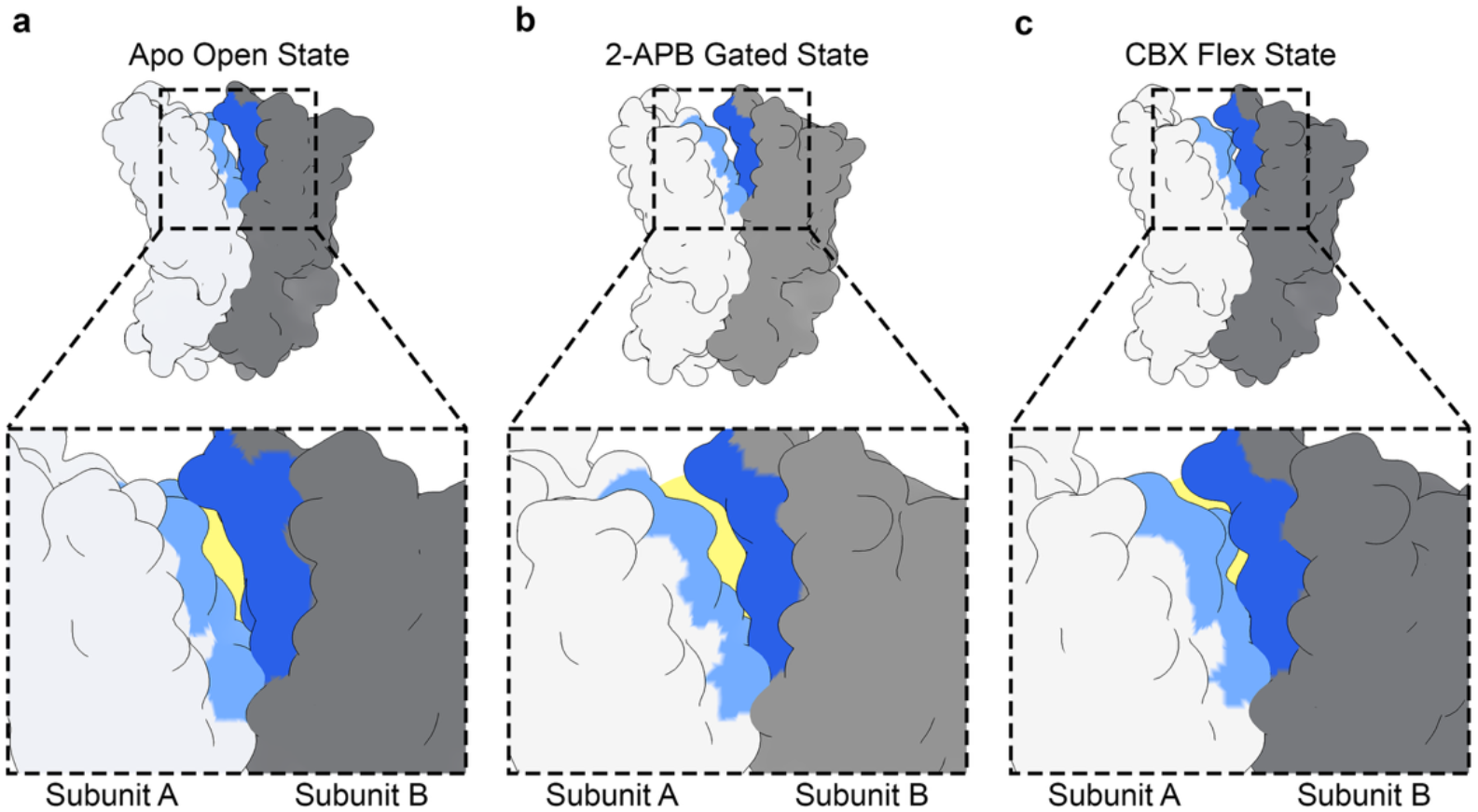
Comparison of the intersubunit fenestration in apo open, 2-APB-gated and CBX-flex states. **a-c**, Surface representations of adjacent connexin subunits showing the intersubunit fenestration in the (*a*) apo open, (*b*) 2-APB-gated and (*c*) CBX-flex states. Subunits A and B are shown in light and dark gray, respectively, with fenestration-lining residues shown in light blue for subunit A and dark blue for subunit B. Insets show enlarged views of the fenestration, with the opening between subunits highlighted in yellow.

---

**Supplementary Table 1:**
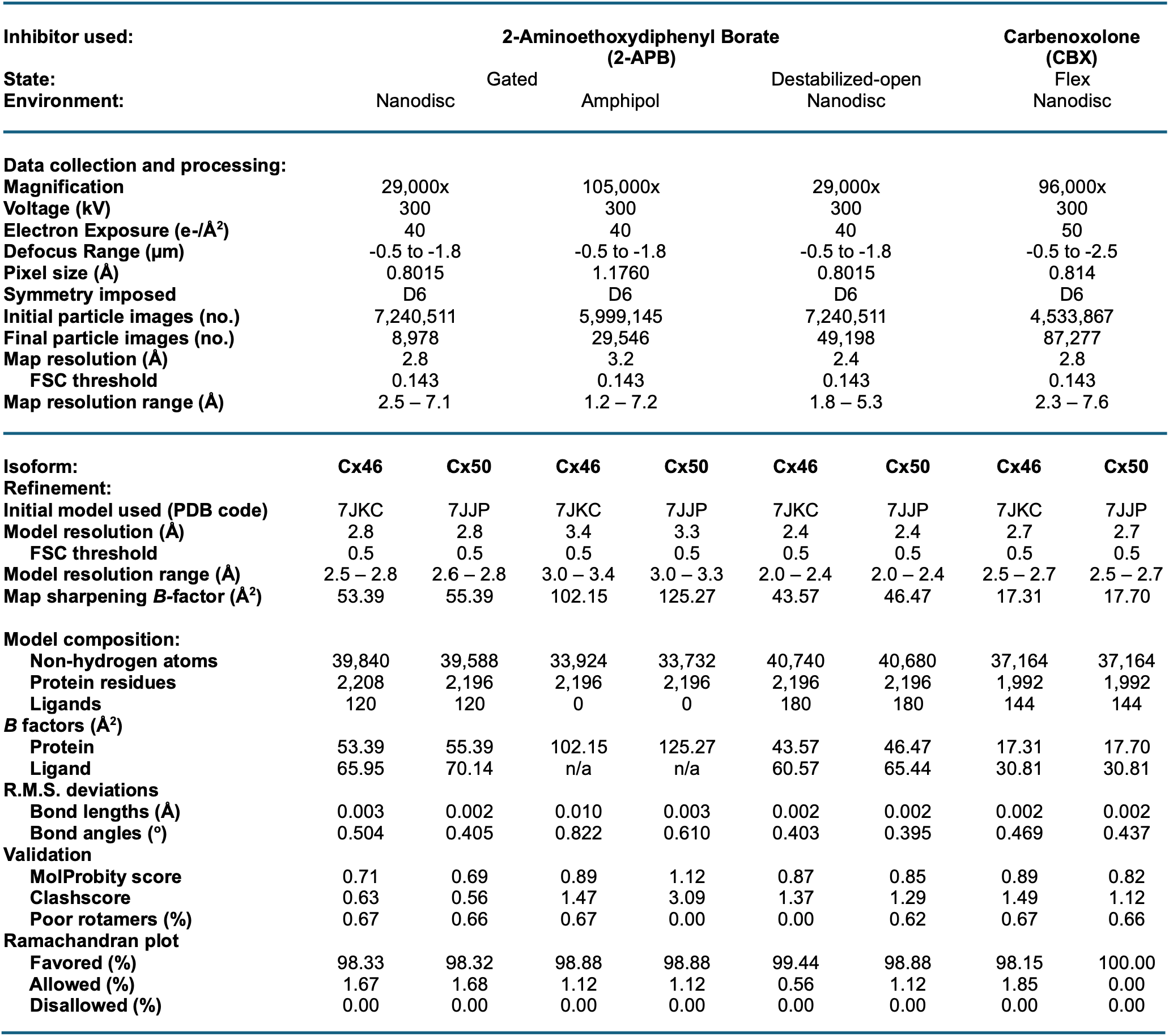
Cryo-EM data collection, refinement and validation statistics.

## SUPPLEMENTARY MOVIE LEGENDS

**Supplemental Movie 1:** 3DVA of Cx46/50 under Apo, 2-APB and CBX conditions.

